# Positive and negative control of helicase recruitment at a bacterial chromosome origin

**DOI:** 10.1101/2021.08.16.456468

**Authors:** Charles Winterhalter, Daniel Stevens, Stepan Fenyk, Simone Pelliciari, Elie Marchand, Nora B Cronin, Panos Soultanas, Tiago R. D. Costa, Aravindan Ilangovan, Heath Murray

## Abstract

The mechanisms responsible for helicase loading during the initiation of chromosome replication in bacteria are unclear. Here we report both a positive and a negative mechanism for directing helicase recruitment in the model organism *Bacillus subtilis*. Systematic mutagenesis of the essential replication initiation gene *dnaD* and characterization of DnaD variants revealed protein interfaces required for interacting with the master initiator DnaA and with a specific single-stranded DNA (ssDNA) sequence located in the chromosome origin (*D*naD *R*ecognition *E*lement, “DRE”). We propose that the location of the DRE within the replication origin orchestrates recruitment of helicase to achieve bidirectional DNA replication. We also report that the developmentally expressed repressor of DNA replication initiation, SirA, acts by blocking the interaction of DnaD with DnaA, thereby inhibiting helicase recruitment to the origin. These findings significantly advance our mechanistic understanding of helicase recruitment and regulation during bacterial DNA replication initiation. Because DnaD is essential for the viability of clinically relevant Gram-positive pathogens, DnaD is an attractive target for drug development.

## INTRODUCTION

Genome replication most often initiates at specific chromosomal loci termed origins. Throughout the domains of life, initiator proteins containing a conserved AAA+ (*A*TPase *A*ssociated with various cellular *A*ctivities) motif assemble at chromosome origins and direct loading of two helicases for bidirectional DNA replication (Bleichert et al., 2017). Interestingly, while the initiation pathway in both bacteria and eukaryotes culminates in ring shaped hexameric helicases encircling a single DNA strand, the molecular mechanisms used to achieve this outcome appear to be distinct (Bell and Kaguni, 2013). Bacteria use their master initiator DnaA to first unwind the chromosome origin (*oriC*) and then load helicases around ssDNA such that they are poised to start unwinding. The eukaryotic initiator ORC (*O*rigin *R*ecognition *C*omplex) also promotes helicase loading, but in this case the annular enzyme is deposited around double-stranded DNA (dsDNA) in a dormant state which must subsequently be activated to form an open complex and encircle a single strand. These distinctions make bacterial DNA replication initiation proteins attractive targets for antibiotic development (Kaguni, 2018; Robinson et al., 2012; van Eijk et al., 2017).

Despite decades of study, the mechanisms underpinning coordinated helicase recruitment and loading to support bidirectional DNA replication initiation in bacteria are unclear (Bell and Kaguni, 2013; Coster and Diffley, 2017; Miller et al., 2019; Ticau et al., 2015). Moreover, bacteria are not known to regulate helicase recruitment, rather they are thought to modulate the onset of DNA replication by controlling the ability of the ubiquitous master initiator DnaA to bind and unwind the chromosome origin.

DnaA is a multifunctional enzyme composed of four distinct domains that act in concert during DNA replication initiation (Fig. S1A) (Messer et al., 1999). Domain IV contains a helix-turn-helix dsDNA binding motif that specifically recognizes 9 base-pair asymmetric sequences called “DnaA-boxes” (consensus 5′-TTATCCACA-3′) (Fujikawa et al., 2003; Fuller et al., 1984; Roth and Messer, 1995).

Domain III is composed of the AAA+ motif that can assemble into an ATP-dependent right-handed helical oligomer (Erzberger et al., 2006; Erzberger et al., 2002; Schaper and Messer, 1997). Domain III also contains the residues required for a DnaA oligomer to interact specifically with a trinucleotide ssDNA binding element termed the “DnaA-trio” (consensus 3′-GAT-5′) (Duderstadt et al., 2011; Ozaki et al., 2008; Richardson et al., 2016). It has been proposed that a DnaA oligomer, guided by DnaA-boxes and DnaA-trios at *oriC*, interacts with one strand of the DNA duplex to promote chromosome origin opening (Duderstadt et al., 2011; Pelliciari et al., 2021; Richardson et al., 2016; Richardson et al., 2019). Additionally, it has been proposed that the AAA+ motif of the DnaA oligomer acts as a docking site for an essential AAA+ helicase chaperone (DnaI in *B. subtilis*, DnaC in *Escherichia coli*), thereby directing the recruitment and correct spatial deposition of helicase onto one DNA strand (Mott et al., 2008).

DnaA domain II tethers domains III/IV to domain I, which acts as an interaction hub. Domain I (DnaA^DI^) facilitates homo-oligomerisation, either directly through a self-interaction (Weigel et al., 1999) or indirectly via accessory proteins such as DiaA and HobA (Keyamura et al., 2007; Natrajan et al., 2009). Domain I also interacts with important regulatory proteins such as HU, Dps and SirA (Chodavarapu et al., 2008a; Chodavarapu et al., 2008b; Jameson et al., 2014; Rahn-Lee et al., 2011) and has weak affinity for ssDNA (Abe et al., 2007). Critically, the most important role of DnaA^DI^ is thought to be recruiting the replicative helicase. This may occur either directly, as for *E. coli* DnaA (Sutton et al., 1998), or indirectly, as for *Bacillus subtilis* DnaA acting as a platform to recruit additional essential replication initiation proteins (Fig. 1A) (Matthews and Simmons, 2018; Smits et al., 2010). Interestingly, in both cases a shared surface on DnaA domain I is suggested to be involved (Fig. S1B-C) (Abe et al., 2007; Keyamura et al., 2009; Martin et al., 2019; Matthews and Simmons, 2018; Seitz et al., 2000).

**Figure 1.**
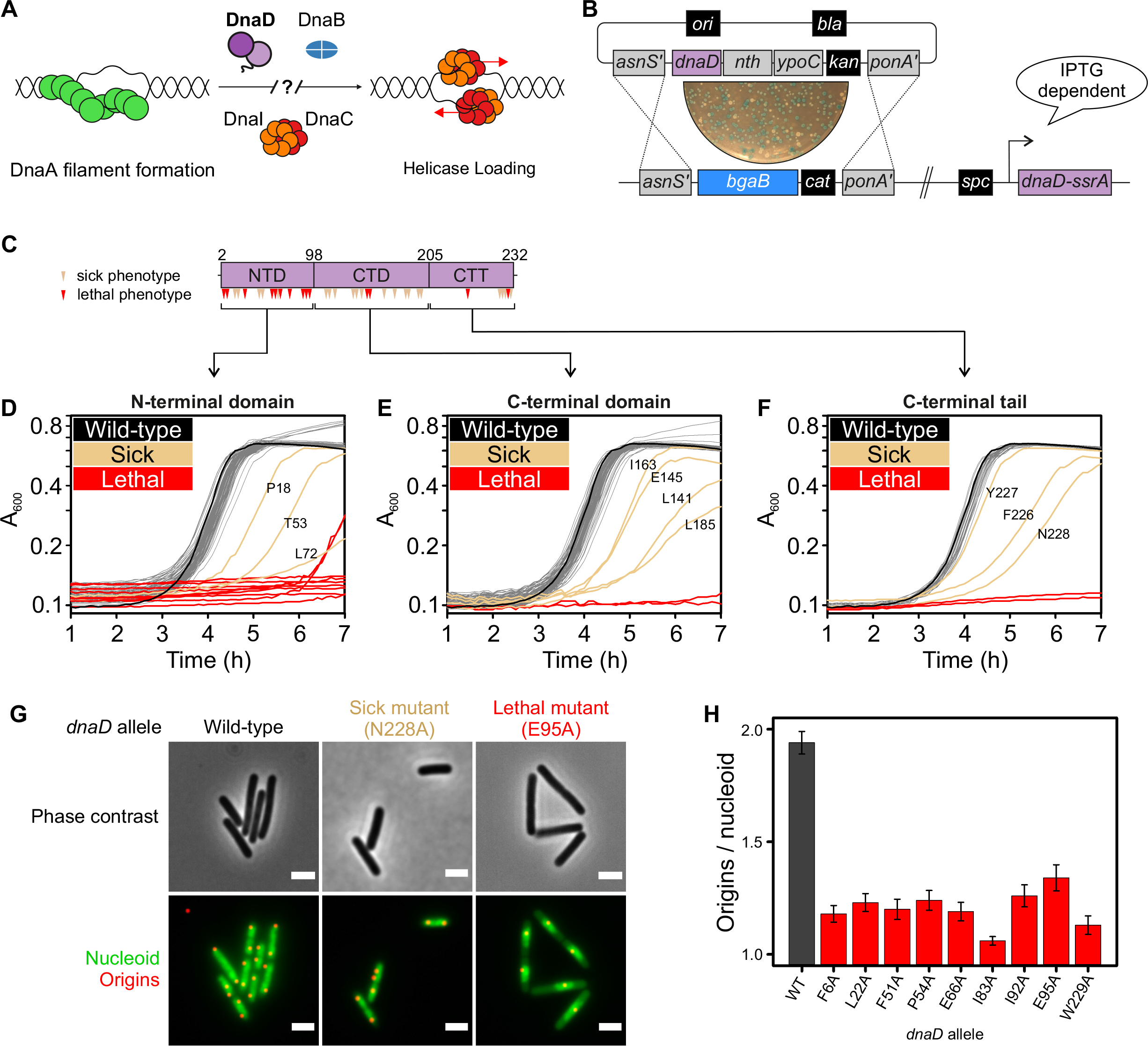
Identification of essential residues in *B. subtilis* DnaD. **(A)** Schematics of the helicase loading pathway in *B. subtilis* showing sequential recruitment of DnaA, DnaD, DnaB and the helicase complex Dnal-DnaC. **(B)** DnaD blue/white screening assay. An integration vector carrying individual *dnaD* substitutions is intergrated by double recombination at the *dnaD* locus, where the native operon has been substitutions is integrated by double recombination at the *dnaD* locu, where the native operson has been replaced by a *bgaB* cassetter allowing screening in the presence of X-gal. Mutant strains harbour the ectopic inducible *dnaD-ssrA* cassette required for viability during transformation and *dnaD* mutant propagation. **(C)** DnaD primary structure. Resuidues marked as red for lethal and beige for altered growth (sporulation defect of slow growth). **(D-F)** Growth analysis of DnaD variants within the N-terminal domain **(D)**, C-terminal domain **(E)** or C-terminal tail **(F)** in the absence of DnaD-SsrA. **(G)** Microscopy analysis of *dnaD* mutants. The Hbs-GFP signal (green) reveals location of the nucleoid within cells, whereas origins are localized by visualizing TetR-mCherry binding a *tetO* array integrated near *oriC* (red). **(H)** Origin to nucleoid ratio for all lethal *dnaD* substitutions where the DnaD variant expression was detectable *in vivo*.

In *B. subtilis,* DnaA recruits DnaD to the chromosome origin and this action is required for the sequential recruitment of DnaB, followed by a complex of the DnaC helicase with its chaperone DnaI (Figure 1A) (Briggs et al., 2012; Marston et al., 2010; Smits et al., 2010). While DnaD and DnaB are known to be essential factors during both DNA replication initiation and restart at repaired replication forks (Bruand et al., 2005), a mechanistic understanding of the activities performed by these replication proteins has remained elusive (Matthews and Simmons, 2018; Rokop and Grossman, 2009; Smits et al., 2010; Smits et al., 2011).

In this paper we focus on DnaD, which will be described as having three domains: N-terminal domain (DnaD^NTD^), C-terminal domain (DnaD^CTD^) and C-terminal tail (DnaD^CTT^) (Fig. 1C). DnaD^NTD^ facilitates oligomerisation (Schneider et al., 2008) and contains a binding site for DnaA (Matthews and Simmons, 2018), while DnaD^CTD^/DnaD^CTT^ is involved in binding DnaA (Martin et al., 2019) and DNA (Carneiro et al., 2006; Huang et al., 2016), as well as untwisting the DNA double helix (Zhang et al., 2006; Zhang et al., 2008).

To explore the role of DnaD in the mechanism of DNA replication initiation, we performed a systematic alanine scan to identify residues essential for DnaD activity within its physiological environment. Structural and functional characterization of DnaD identified regions required for protein:protein and protein:DNA interactions. The results suggest that DnaD is recruited to a specific strand of the open complex formed at *oriC* via a new ssDNA binding motif (the DRE), thus providing a potential route for directing helicase loading. Moreover, we find that the recruitment of DnaD to DnaA in *B. subtilis* is developmentally regulated by SirA.

## RESULTS

DnaD is an essential DNA replication initiation protein in the model organism *B. subtilis* and in opportunistic pathogens such as *Staphylococcus aureus* and *Streptococcus pneumoniae* (Chaudhuri et al., 2009; Kobayashi et al., 2003; Liu et al., 2017). However, the activities required for DnaD to perform its role at the chromosome origin *in vivo*, and the mechanisms underlying those activities, are unclear. To address these questions, we sought to identify essential amino acids in *B. subtilis* DnaD and then to determine the function of each essential residue.

### Identification of essential residues in DnaD necessary for cellular DNA replication initiation

Functional analysis of bacterial DNA replication initiation proteins *in vivo* is challenging because they are required for viability; mutation of an essential feature will be lethal, while mutations that severely disable function can result in the rapid accumulation of compensatory suppressors. To circumvent these issues, a bespoke inducible complementation system was developed for *dnaD* (*PHAT dnaD-ssrA*, Fig. S2 and Supplementary text). Upon repression of the ectopic *dnaD-ssrA*, the functionality of *dnaD* alleles at the endogenous locus can be determined.

A plasmid for allelic exchange of the endogenous *dnaD* gene was created (Figure S3). Using this as a template, a library of 222 single alanine substitution mutants (all codons save for the start, stop and naturally occurring alanine) was generated and sequenced. To ensure mutagenesis of the native *dnaD* following transformation, a recipient strain was constructed containing both the *dnaD* operon replaced by *bgaB* (encoding the enzyme β- galactosidase) and the inducible *dnaD* complementation system (Fig. S3). Thus, replacement of *bgaB* with *dnaD* alleles can be detected on selective media supplemented with a chromogenic substrate (white colonies, Fig. 1B) and confirmed by chloramphenicol sensitivity.

Following construction of the *dnaD* alanine substitution library, strains were grown in a plate reader to assess the functionality of each mutant. The data revealed growth defects for several alanine substitutions, spread throughout the protein (Fig. 1C-F). Immunoblots were used to determine whether the DnaD variants were being stably expressed (Fig. S4A, C, E, G), and for those with detectable levels of protein a spot-titre assay was used to confirm growth phenotypes (Fig. S4B, D, F). We note that the level of DnaD sufficient to sustain colony formation is below the detection level of our immunoblots (Fig. S5). However, for the least ambiguous interpretation of the results, we focussed on essential alanine substitutions that were expressed near the wild-type level.

While essential for DNA replication initiation in *B. subtilis*, DnaD has also been implicated in other key cellular processes including chromosome organization and DNA repair (Collier et al., 2012; Ishikawa et al., 2007; Smits et al., 2011; Zhang et al., 2005). To ascertain whether *dnaD* alanine mutants were specifically impaired in DNA replication initiation, we further characterized chromosome content in these strains using fluorescence microscopy.

During slow steady-state growth, wild-type *B. subtilis* cells typically display a pair of chromosome origins per nucleoid, each orientated towards a cell pole (Fig. 1G) (Webb et al., 1997). In contrast, when chromosome replication is inhibited, nucleoids typically contain a single *oriC* signal located near the centre of the bulk DNA (Imai et al., 2000). Therefore, a strain harbouring *hbs-gfp* to detect the nucleoid (Kohler and Marahiel, 1997) and a fluorescent reporter-operator system (*tetO* array with *tetR-mCherry*) (Wang et al., 2014) to detect the chromosome origin region was used to evaluate the impact of *dnaD* mutants on DNA replication (Fig. S6A). Strains were imaged following repression of the ectopic *dnaD-ssrA* for 90 minutes. All *dnaD* alanine mutants produced a phenotype characteristic of non-replicating chromosomes, with well separated chromosomes often containing a single TetR-mCherry focus (Fig. 1G-H and S6B). Taken together, this analysis identified 14 alanine substitutions in DnaD that retained detectable protein expression and produced a growth phenotype, nine of which were essential for DNA replication initiation *in vivo* (Fig. 1E-G).

### A DnaD tetramer is necessary for DNA replication initiation in vivo

The crystal structure of DnaD^NTD^ was solved as a symmetric homodimer, while biochemical experiments and structural modelling suggest assembly into a tetramer or higher-order oligomer (Briggs et al., 2012; Schneider et al., 2008). However, the active quaternary structure of the protein *in vivo* was not known. The DnaD alanine scan showed that replacement of either Phe6 or Leu22 was lethal (Fig. S4B-C). Mapping these residues onto the DnaD^NTD^ crystal structure (Fig. 2A) reveals that Leu22 is buried within the proposed dimerization interface and that Phe6 is exposed towards a predicted dimer:dimer interface (see Fig. S7A, which includes the positions of expressed alanine substitutions with growth defects and the unexpressed DnaD^K3A^ (Fig. S4A). Therefore, we investigated DnaD^F6A^ and DnaD^L22A^ for defects in oligomerisation.

**Figure 2.**
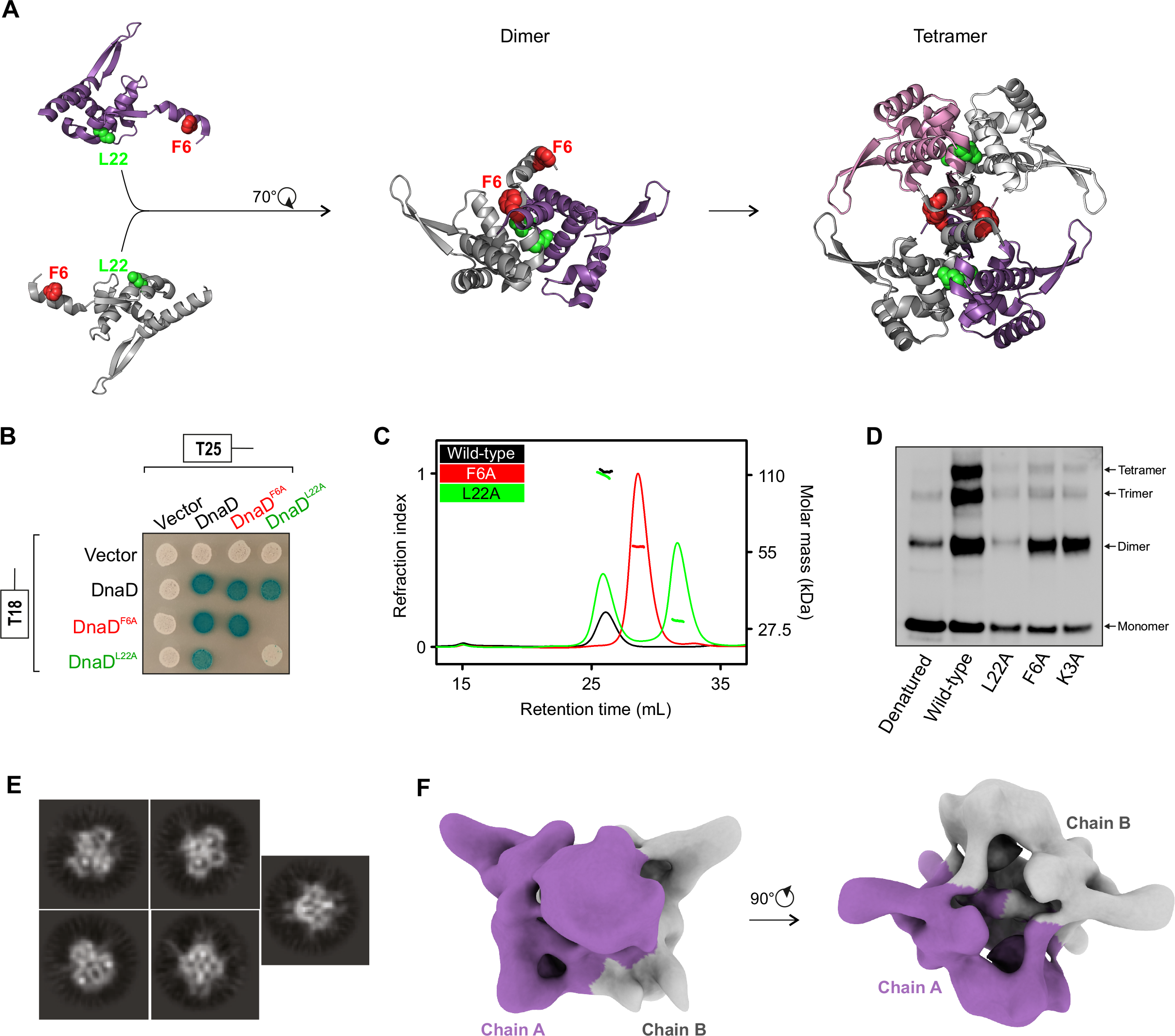
Lethal alanine substitutions in DnaD disrupt tetramer formation. **(A)** Schematics of DnaD N-terminal domain oligomerisation pathway involving key substitutions F6A and L22A. **(B)** Bacterial two-hybrid assay showing the effect of mutants DnaD^L22A^ and DnaD^F6A^ on self-interaction. **(C)** SEC-MALS analysis of DnaD variants. The UV spectrum was normalised as a refraction index and molar mass corresponding to each protein represented as a dots overlapping the peaks. **(D)** Immunoblot following migration and transfer of BS^3^ crosslinked DnaD species using SDS-PAGE. **(E)** 2D classes observed by cryo-EM. **(F)** Cryo-EM map of a DnaD dimer.

To begin assessing the DnaD self-interaction a bacterial two-hybrid assay was employed. Full-length *dnaD* alleles were fused to catalytically complementary fragments of the *Bordetella pertussis* adenylate cyclase (T25 and T18)(Karimova et al., 1998). Two-hybrid analysis showed that wild-type DnaD and DnaD^F6A^ self-interact, whereas DnaD^L22A^ lost this capability (Fig. 2B). All DnaD proteins reported a positive interaction with wild-type DnaD, indicating that all *dnaD* alleles were being functionally expressed in the heterologous host (Fig. 2B). These results suggest that Leu22 is involved in DnaD dimer formation.

To further interrogate the quaternary structure of DnaD, we purified DnaD^L22A^ and DnaD^F6A^ and characterised these variants by size exclusion chromatography (SEC) (Hagel, 2001) followed by multiple angle light scattering (MALS) (Wyatt, 1993). Wild-type DnaD was observed to run as a stable tetramer of approximately 113 kDa (theoretical molecular weight of 110 kDa) (Fig. 2C). SEC-MALS analysis showed that >50% of DnaD^L22A^ dissociated into a 29 kDa monomer, whereas DnaD^F6A^ was eluted exclusively as 57 kDa species, consistent with the protein forming a stable dimer (Fig. 2C). Crosslinking with amine-specific bis(sulfosuccinimidyl)suberate (BS^3^) confirmed that DnaD^F6A^ was competent to form a dimer but defective to form a tetramer (Fig. 2D). Returning to the *dnaD* alanine scan we appreciated that *dnaD^K3A^* was also lethal, albeit poorly expressed *in vivo* (Fig. S4A). Crosslinking showed that DnaD^K3A^ could also form a dimer but not a tetramer, akin to DnaD^F6A^ (Fig. 2D). Taken together, the data indicate that DnaD tetramerization is mediated by residues located near the N-terminus and that adopting this quaternary state is necessary to support DNA replication initiation *in vivo*.

### Architecture of a DnaD dimer determined by cryo electron microscopy

To elucidate the quaternary structure of DnaD, we characterized the structure of the full-length protein using single particle cryo electron microscopy (cryo-EM). Although the wild-type tetrameric DnaD was used, it was clear from the cryo-EM data that only a pair of proteins was observable (Fig. 2E). Data analysis from 2D classes (Fig. 2E) and image processing revealed a 10 Å resolution map of a DnaD dimer (Fig. 2F and S7B). DnaD^CTD^ subunits and the β-hairpin within the DnaD^NTD^ were immediately identified within the cryo-EM map, and a poly alanine model of full-length DnaD encompassing a pair of DnaD^NTD^ and DnaD^CTD^ could be recognised (Fig. S7C). While the previously published DnaD^CTD^ structure (PDB 2zc2) agrees well with the cryo-EM model, some differences were observed with the arrangement of α-helices and β-strands described in the crystal structure of the DnaD^NTD^ (PDB 2v79) (Schneider et al., 2008). The conditions surrounding these two states and their functional relevance were not explored further. These differences notwithstanding, the cryo-EM map reveals for the first time that the DnaD^NTD^ and DnaD^CTD^ pack against each other to form a compact structure. The placement of the DnaD^CTD^ suggest that the DnaD^CTT^, which was not assigned within the map, would extend from a location on the opposite face to the proposed dimer:dimer interface (Fig. S7C). Implications of the DnaD structure on protein function are explored below.

### The interaction between DnaD^CTT^ and DnaB is necessary for DNA replication initiation in vivo

The alanine scan indicated that a cluster of residues in the unstructured C-terminal tail of DnaD were critical for cell growth, particularly Trp229 which is essential (Fig. 3A and S4F-G). Phylogenetic analysis indicated that Trp229 was conserved in species harbouring both *dnaD* and *dnaB*, but not *dnaD* alone, suggesting that the DnaD^CTT^ could be an interaction site for DnaB (Fig. S8A).

**Figure 3.**
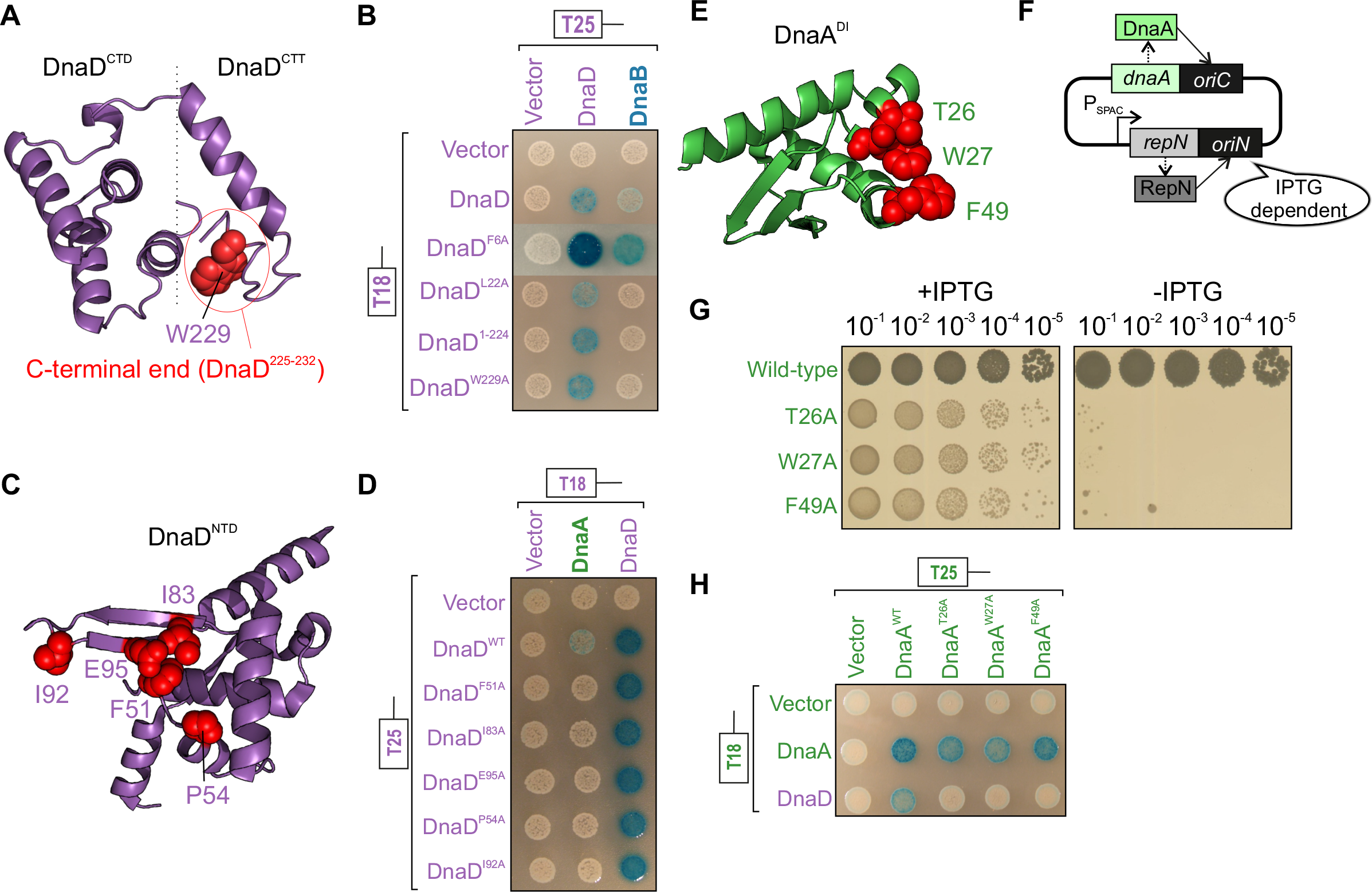
Lethal alanine substitutions in DnaD disrupt interactions with DnaA and DnaB. **(A)** Essential residues of DnaD, required for the interaction with DnaA, mapped onto the DnaD^NTD^ crystal structure (PDB 2V79). **(B)** Bacterial two-hybrid assay showing loss of interaction between DnaD^NTD^ variants and DnaA in the context of full-length proteins. **(C)** Essential residues of DnaD, required for the interaction with DnaB, mapped onto a DnaD^CTD/CTT^ model. **(D)** Bacterial two-hybrid assay showing loss of interaction between DnaD variants and DnaB in the context of full-length proteins. **(E)** Essential residues of DnaA, required for the interaction with DnaD, mapped onto the DnaA^DI^ crystal structure (PDB 4TPS). **(F)** Schematics of the inducible *repN/oriN* system used to bypass mutations affecting DnaA activity in *B. subtilis*. **(G)** Spot-titre analysis of DnaA^DI^ variants using the inducible *oriN* strain. **(H)** Bacterial two-hybrid assay showing loss of interaction between DnaA variants and DnaD in the context of full-length proteins.

To test this hypothesis, two-hybrid analysis was used to probe for a direct protein:protein interaction. The results showed that wild-type DnaD and DnaB interact and that both the lethal allele *dnaD^W229A^* and deletion of the last eight amino acids break this interaction (Fig. 3B). It was also observed that the monomeric DnaD^L22A^ was unable to interact with DnaB, whereas the dimeric DnaD^F6A^ retained this capability (Fig. 3B). All protein variants retained a self-interaction with wild-type DnaD, showing that they were being functionally expressed (Fig. 3B). These results indicate that the interface between the distal end of DnaD^CTT^ and DnaB is essential for DNA replication initiation *in vivo*, and they suggest that DnaB recognition requires DnaD assembly into a homodimer.

Previous studies using protein truncation variants indicated that the DnaD^NTD^ interacts with DnaB (Matthews and Simmons, 2018). The observation that mutations in the DnaD^CTT^ abolish the interaction with DnaB (where the DnaD^NTD^ is present) suggests that different interactions are being detected in these assays. We note that the N-terminal domains of DnaD and DnaB share structural homology (Fig. S8B) and both promote dimerization/tetramerization, such that the truncated variants may be able to interact differentially in a two-hybrid experiment.

### The interaction between DnaD^NTD^ and DnaA is necessary for DNA replication initiation in vivo

Models for the interaction between DnaD and DnaA have been proposed based on binding experiments using truncated proteins. These studies indicated that residues in the DnaD^NTD^ (Matthews and Simmons, 2018) and the DnaD^CTD^ (Martin et al., 2019) each contributed to DnaA binding. From the alanine scan it was observed that three of the proposed residues at the DnaA interface of the DnaD^NTD^ are essential (Phe51, Ile83, Glu95). Additionally, we identified two other lethal substitutions (DnaD^P54A^ and DnaD^I92A^) that mapped near these sites on the structure, suggesting they could also be involved in the DnaA interface (Fig. 3C and S4D-E).

In contrast, none of the residues in the DnaD^CTD^ were found to be essential (Fig. S9A-C). To investigate whether the interface between DnaD^CTD^ and DnaA was robust and single alanine substitutions were insufficient to disrupt binding, DnaD variants encoding multiple alanine substitutions were constructed (*dnaD^L129A/I132A^, dnaD^Y130A/E134A/E135A^, dnaD^I132A/E134A/E135A^, dnaD^K164A/K168A/E169A/V171A^*) (Fig. S9D-G). However, all of these *dnaD* alleles were viable (Fig. S9C). These results indicate that the interface between DnaA and the DnaD^CTD^ is not essential for DNA replication initiation *in vivo*. The DnaD^CTD^-DnaA interaction could play an auxiliary role that assists DnaD binding DnaA, or alternatively it could become important during certain environmental conditions or cell stresses.

Two-hybrid analysis was used to investigate whether alanine substitutions in DnaD^NTD^ perturb the interaction with DnaA. It was known that expression of *B. subtilis* DnaA in *E. coli* perturbs DNA replication and inhibits cell growth, presumably by competing with the endogenous homolog for binding DnaA-boxes within *oriC* (Andrup et al., 1988; Krause and Messer, 1999), and it was previously reported that interactions between full-length *B. subtilis* DNA replication initiation proteins could not be detected (Matthews and Simmons, 2018). To circumvent the toxicity elicited by *B. subtilis* DnaA, we reduced selective pressure by constructing a derivative of the *E. coli* two-hybrid strain with a deletion of the *rnhA* gene. This strain can initiate replication at stable R-loops that are normally removed by RNase HI and this mode of DNA replication initiation is independent of DnaA and *oriC* (Kogoma and von Meyenburg, 1983). Using this approach an interaction between the full-length DnaD and DnaA proteins was detected (Fig. 3D). In contrast, alanine substitutions within the proposed DnaD^NTD^ interface for DnaA abrogated this association (Fig. 3D). All DnaD^NTD^ variants retained the ability to self-interact, indicating that they were being functionally expressed. These results indicate that a direct interaction between DnaA and DnaD^NTD^ is essential for DNA replication initiation *in vivo*.

### The interaction between DnaA^DI^ and DnaD is necessary for DNA replication initiation in vivo

Having identified sites on DnaD for protein:protein interactions, we further investigated the complex formed with the master initiator DnaA. Previous two-hybrid and NMR studies identified residues on the surface of DnaA^DI^ that interact with DnaD (Martin et al., 2019; Matthews and Simmons, 2018). To investigate the physiological relevance of the proposed DnaA^DI^ interface with DnaD *in vivo*, we replaced the endogenous *dnaA* gene with mutant variants encoding alanine substitutions at key residues (*dnaA^T26A^*, *dnaA^W27A^*, *dnaA^F49A^*) (Fig. 3E). To enable identification of essential amino acid residues without selecting for suppressor mutations, we utilized a strain in which DNA replication can initiate from a plasmid origin *(oriN*) integrated into the chromosome (Fig. 3F) (Richardson et al., 2016). Activity of *oriN* requires its cognate initiator protein RepN; both factors act independently of *oriC*/DnaA (note the RepN/*oriN* system does require DnaD and DnaB for function) (Fig. S10A-C) (Hassan et al., 1997). Expression of *repN* was placed under the control of an IPTG-inducible promoter, thus permitting both the introduction of mutations into *dnaA* and their subsequent analysis following removal of the inducer to repress *oriN* activity. Cultures were grown overnight in the presence of IPTG and then serially diluted onto solid media. The results showed that the *dnaA^T26A^*, *dnaA^W27A^* and *dnaA^F49A^* mutants all inhibited colony formation (Fig. 3G). Immunoblot analysis indicated that all of the DnaA variants were expressed at a level similar to wild-type (Fig. S10B). This analysis indicates that residues Thr26, Trp27 and Phe49 are essential for DnaA activity *in vivo*.

Two-hybrid analysis confirmed that alanine substitutions in DnaA at either Thr26, Trp27 or Phe49 inhibit the interaction with DnaD (Fig. 3H). These DnaA variants retained the ability to self-interact with the wild-type protein, indicating that they are being functionally expressed (Fig. 3H). Therefore, the essential residues in DnaA^DI^ are required to bind DnaD. To investigate whether DnaA^DI^ and DnaD^NTD^ were sufficient to form a complex, pull-down assays between protein domains His6-DnaA^DI^ and DnaD^NTD^ were performed (Fig. S11A). Following expression of His6-DnaA^DI^ and DnaD^NTD^ in *E. coli*, cells were lysed and His6-DnaA^DI^ was captured using an immobilized nickel affinity chromatography spin column. While the wild-type DnaD^NTD^ was able to bind wild-type His6-DnaA^DI^, amino acid substitutions in either protein domain greatly reduced the retention of DnaD^NTD^ (Fig. S11B). Staining of SDS-PAGE revealed that all protein domains were being overexpressed to similar levels (Fig. S11C) and immunoblotting confirmed the identity of each polypeptide (Fig. S11D). These studies support and extend the previously proposed model for DnaA^DI^ interacting with DnaD^NTD^ (Matthews and Simmons, 2018), critically showing that this protein:protein interface is essential for DNA replication initiation *in vivo*.

### The interaction of DnaA^DI^ with DnaD^NTD^ is required to recruit DnaD to the chromosome origin

It has been observed that DnaA recruits DnaD to the replication origin (Smits et al., 2010) and we hypothesized that this could be the essential function of the DnaA^DI^-DnaD^NTD^ interaction. To test this model, we employed chromatin immunoprecipitation (ChIP). To support growth of lethal *dnaA* mutants, a strain harbouring a constitutively active version of *oriN* in the chromosome was used. In all cases, DnaA^DI^ variants remained specifically enriched at *oriC* while recruitment of DnaD was abolished (Fig. 4A). Note that in these strains DnaD remained enriched at *oriN,* as expected (Fig. 4B) (Smits et al., 2011). These data are consistent with the proposal that an essential function of the DnaA^DI^-DnaD^NTD^ interaction is to recruit DnaD to the chromosome origin. Intriguingly, the surface of DnaA^DI^ interacting with DnaD is also the binding site for the developmentally expressed DNA replication inhibitor SirA, raising the possibility that SirA could compete with DnaD for binding DnaA (Fig. 4C) (Jameson et al., 2014; Rahn-Lee et al., 2011).

**Figure 4.**
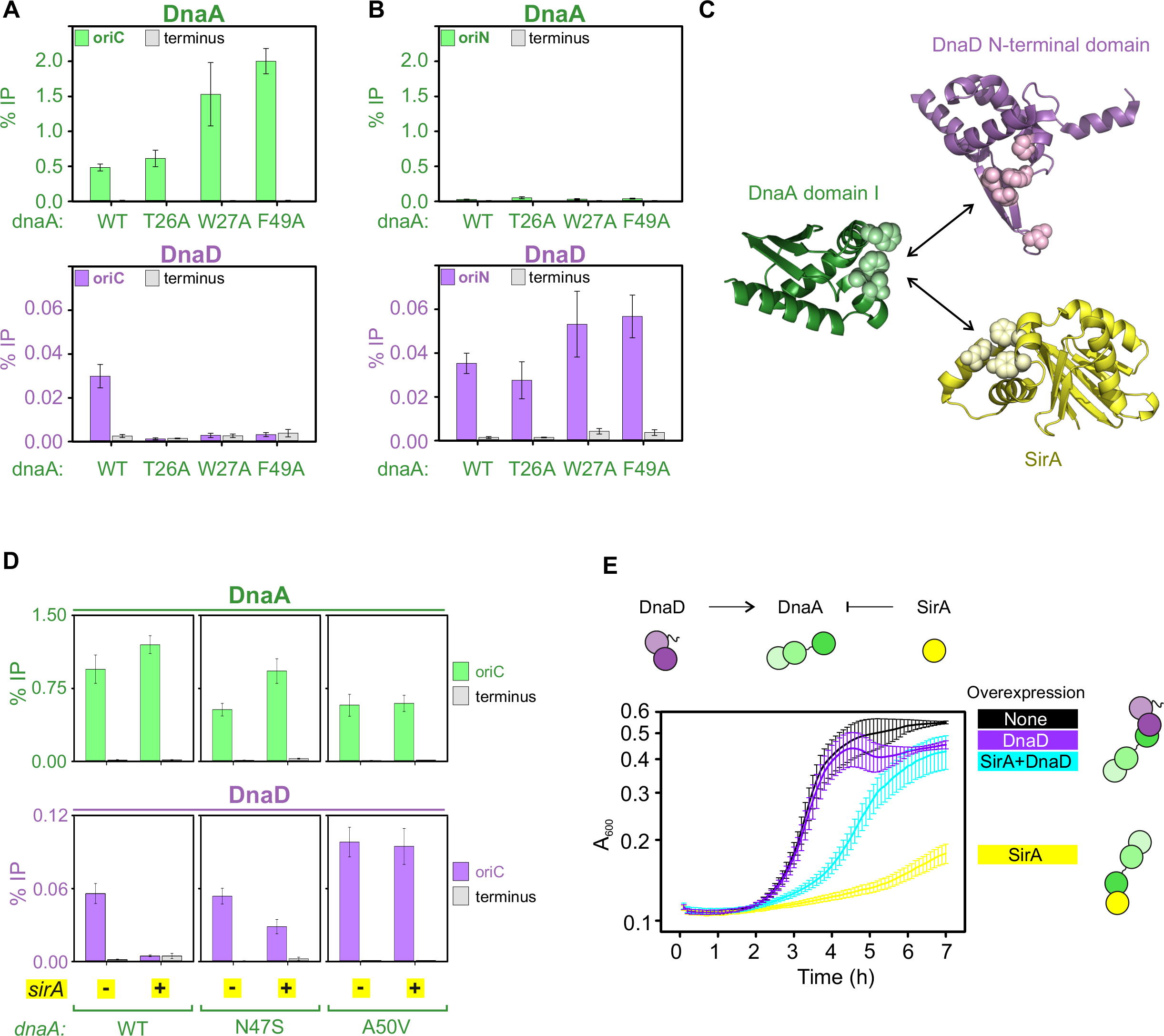
SirA binds to DnaA^DI^ and directly inhibits DnaD recruitment to *oriC*. **(A)** ChIP of DnaA proteins (wild-type, T26A, W27A and F49A) and DnaD at *oriC*. Primers used for the origin anneal within the *incC* region. **(B)** ChIP of DnaA proteins (wild-type, T26A, W27A and F49A) and DnaD at *oriN*. **(C)** Crystal structures of DnaD^NTD^ (PDB 2V79), DnaA^DI^ and SirA (PDB 4TPS) highlighting residues at the protein:protein interfaces. **(D)** ChIP of DnaA and DnaD at *oriC* following overexpression of SirA. **(E)** Growth assay overexpressing SirA with and without DnaD overexpression. None indicates no inducer, DnaD overexpression with 0.35% xylose, SirA overexpression with 0.035 mM IPTG, DnaD and SirA simultaneous overexpression with 0.35% xylose and 0.035 mM IPTG. Error bars indicate the standard error of the mean for at least 3 biological replicates.

### SirA binding to DnaA inhibits recruitment of DnaD and DnaB to oriC

During endospore development *B. subtilis* requires two chromosomes, one for each differentiated cell type (Errington, 2003). To help ensure diploidy after executing the commitment to sporulate, cells express the negative regulator of DNA replication initiation SirA (Rahn-Lee et al., 2009; Wagner et al., 2009). It was proposed that SirA represses DNA replication initiation by inhibiting DnaA binding to *oriC* (Rahn-Lee et al., 2011). However, in the previous study SirA activity was analysed following artificial activation of sporulation, a complex developmental pathway involving the activation and/or induction of hundreds of genes including the master regulator Spo0A (Fawcett et al., 2000), which itself is known to inhibit DNA replication initiation by binding at *oriC* (Boonstra et al., 2013). Therefore, considering the data presented above, we hypothesized that SirA might occlude DnaD from binding to DnaA, thereby inhibiting recruitment of DnaD to *oriC*.

Using a strain containing *sirA* under the control of an IPTG-inducible promoter, SirA was expressed for 30 minutes during mid-exponential growth. ChIP of wild-type DnaA revealed stable enrichment at *oriC* following SirA expression (Fig. 4D), indicating that under these conditions SirA does not inhibit DnaA binding to DNA. ChIP of DnaD showed enrichment was abolished, consistent with the model that SirA inhibits DnaD binding to DnaA. Furthermore, enrichment of the helicase loader DnaB at *oriC*, which requires prior binding of DnaD (Smits et al., 2010), was also lost following induction of *sirA* (Fig. S12). When the ChIP experiments were repeated using alleles of *dnaA* (N47S, A50V) that suppress SirA by inhibiting its binding to DnaA^DI^ (Rahn-Lee et al., 2011), enrichment of DnaD at *oriC* was restored to a degree that correlated with the penetrance of the *dnaA* suppressor mutations (Fig. 4D) (Jameson et al., 2014).

To investigate whether SirA and DnaD binding to DnaA^DI^ is mutually exclusive, we set up a competition experiment. A strain was engineered with ectopic copies of *sirA* and *dnaD* under the control of inducible promoters (IPTG and xylose, respectively; Fig. S13A). While expression of SirA alone inhibited growth, co-expression with DnaD significantly ameliorated this effect (Fig. 4E and S13B-D). However, expression of DnaD variants defective for binding DnaA (DnaD^F51A^, DnaD^I83A^ or DnaD^E95A^) did not reverse the SirA-mediated growth inhibition (Fig. S13E-F). Taken together, the results suggest that SirA inhibits DNA replication initiation by directly occluding the binding of DnaD to DnaA.

### DnaD^CTT^ ssDNA binding activity is essential for DNA replication initiation in vivo

It has been established that DnaD has an affinity for DNA and previous studies employing protein deletions reported that this activity involves the DnaD^CTT^ (Huang et al., 2016; Marston et al., 2010; Smits et al., 2011). Therefore, it was conspicuous that the alanine scan did not identify an essential residue suggesting a role in DNA binding within this protein domain (Fig. 1C). Alignment of DnaD^CTT^ homologues showed the recurrence of positively charged and aromatic residues within this domain (Fig. 5A). Based on these findings, we hypothesized that the DnaD^CTT^ contains a robust DNA binding motif.

**Figure 5.**
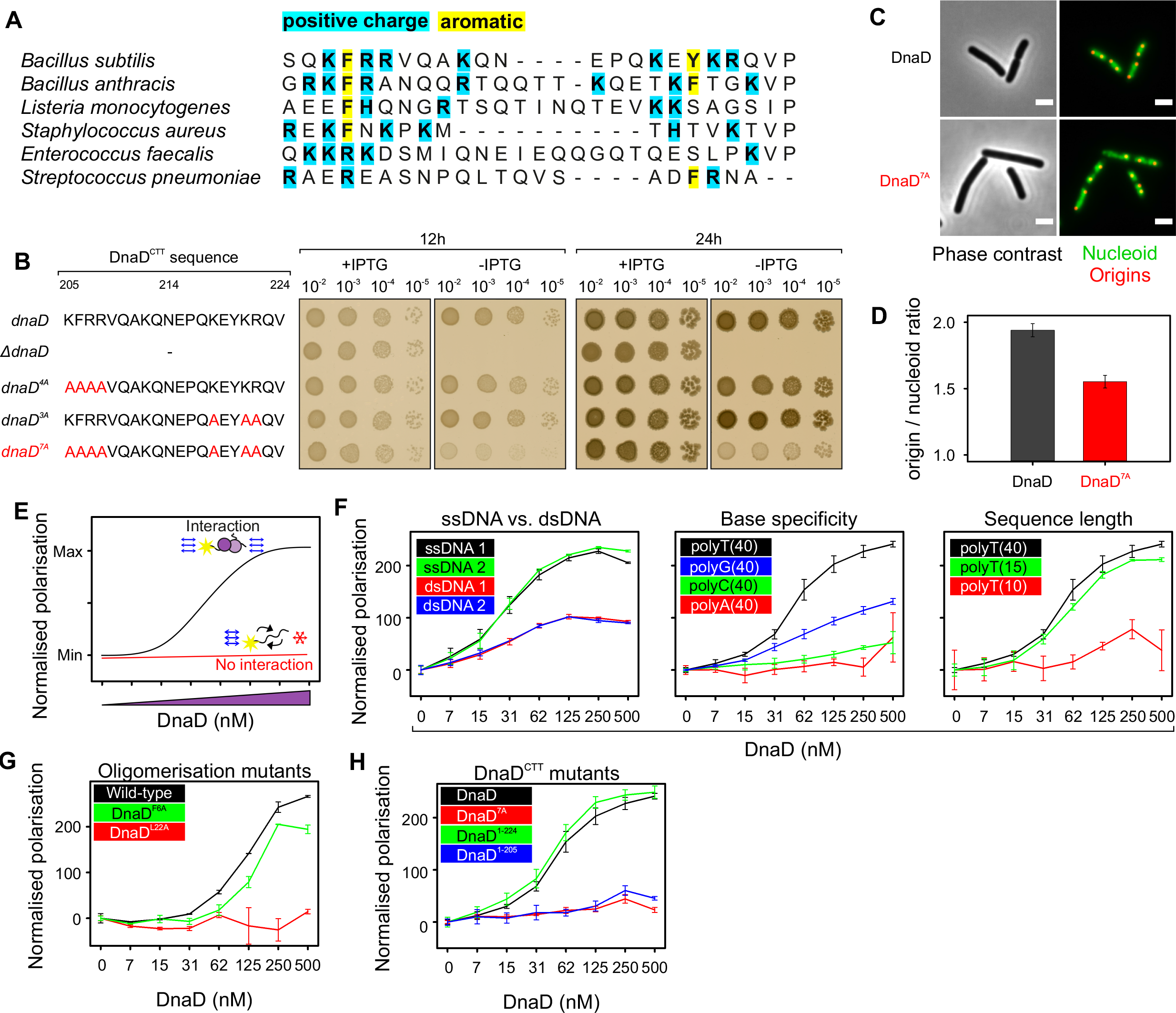
Alanine substitutions in the DnaD C-terminal tail inhibit DNA replication initiation *in vivo* and ssDNA binding *in vitro*. **(A)** Alignment of DnaD homologs showing the recurrence of positively charged and aromatic residues within the DnaD^CTT^. **(B)** Spot-titre analysis of multiple alanine substitutions om the DnaD^CTT^. **(C)** Fluorescence microscopy showing altered nucleoid (*hbs-gfp*, green) and origins of the chromosome (*tetR-mCherrry* bound to a *tetO* array, red) in the DnaD^7A^ mutant. **(D)** Quantification of origins per nucloid in the DnaD^7A^ strain. **(E)** Illustration of the fluorescence polarisation assay. **(F)** Fluorescence polarization analysis of wild-type DnaD binding to a range of DNA substrates. **(G)** Fluorescence polarization analysis of the oligomerisation mutant DnaD^L22A^ (monomer) with ssDNA. **(H)** Fluorescence polarization analysis of variants within the DnaD^CTT^.

*dnaD* alleles encoding multiple alanine substitutions with the DnaD^CTT^ were constructed. Spot titre analysis revealed that substitutions replacing two clusters of residues (DnaD^7A^) resulted in an observable growth defect (Fig. 5B) and immunoblots confirmed the expression of each protein (Fig. S14A). Characterisation of DnaD^7A^ by fluorescence microscopy revealed an apparent DNA replication defect, with cells containing a lower number of *oriC* per nucleoid (Fig. 5C-D). Marker frequency analysis confirmed that DnaD^7A^ displays a decreased DNA replication initiation frequency compared to wild-type cells (Fig. S14B).

To directly investigate the DNA binding activity of DnaD *in vitro*, we established a fluorescence polarization assay to detect the binding of purified DnaD to fluorescein labelled substrates (Fig. 5E) (Moerke, 2009). It was found that wild-type DnaD binds ssDNA with a higher affinity than dsDNA, it displays a preference for thymidine, and it requires a substrate at least 15 nucleotides long (Fig. 5F). Moreover, using oligomerization mutants it was observed that monomeric DnaD^L22A^ could not bind ssDNA, whereas dimeric DnaD^F6A^ retained this activity with a binding profile comparable to wild-type (Fig. 5G).

Next we purified several DnaD variants with alterations to the C-terminal tail, including DnaD^7A^ and two truncations, which removed either the DnaB interaction patch (DnaD^1-224^) or the entire C-terminal tail containing the putative ssDNA binding region (DnaD^1-205^). Both DnaD^7A^ and DnaD^1-205^ were unable to interact with a fluorescently labelled polythymidine (dT40) substrate, whereas DnaD^1-224^ retained this activity (Fig. 5H). Size exclusion chromatography showed that the both DnaD^7A^ and DnaD^1-205^ assembled into a tetramer (Fig. S14C-D), indicating that the overall structure of the proteins remained intact. Combined with the *in vivo* analysis, the results suggest that the essential activity of DnaD located between residues 205-224 is to bind ssDNA.

### DnaD recognizes a specific single-strand DNA binding element within the unwinding region of oriC

During the interrogation of DnaD ssDNA binding activity *in vitro,* we found that the wild-type protein had the highest affinity for a substrate with a sequence found within the unwinding region of *B. subtilis oriC*, the complement of the DnaA-trios (5’-CTACTATTACTTCTACTA-3’) (Fig. 6A-B). Based on this sequence, and on the observation that DnaD binds the dT18 substrate better than other homopolymeric ssDNA (Fig. S15A-B), we hypothesized that thymidine might be a specificity determinant.

**Figure 6.**
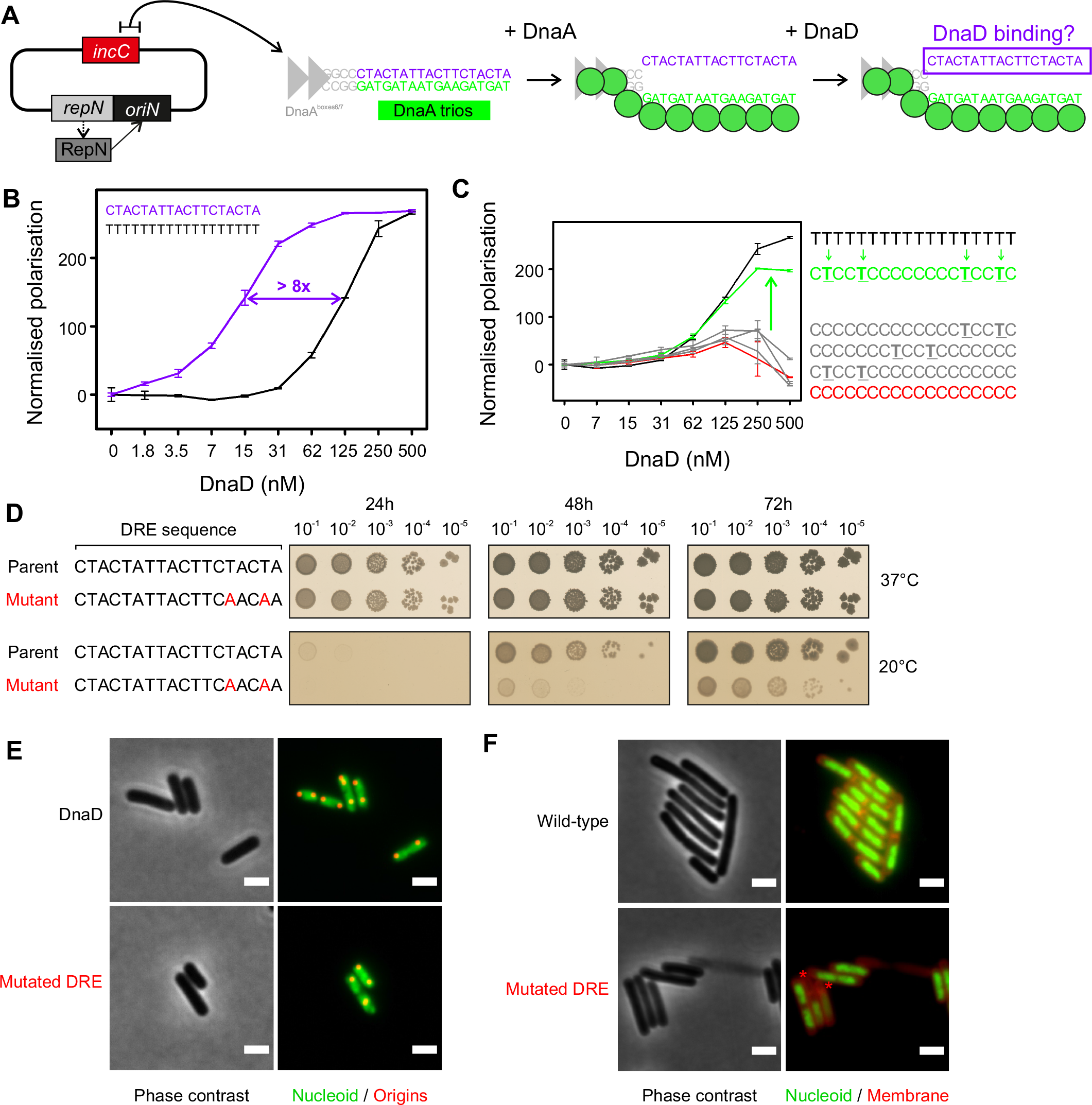
The DRE promotes specific ssDNA binding activity of DnaD *in vitro* and is required for efficient DNA replication initiation *in vivo*. **(A)** Illustration of the proposed basal origin unwinding mechanism involving DnaA oligomer formation on DnaA-trios. **(B)** Fluorescence polarisation analysis of wild-type DnaD on native and non-native ssDNA substrates. **(C)** Fluorescence polarisation analysis of wild-type DnaD on non-specific ssDNA backbones containing various 5’-TnnT-3’ repeats . **(D)** Spot titre analysis of the DRE mutant (5’-TnnT-3’ to 5’-AnnA-3’). **(E)** Fluorescence microscopy showing altered nucleoid (Hbs-GFP, green) and origins of the chromosome (TetR-mCherry bound to a *tetO* array, red) in the DRE mutant shown in (D) when cells were grown at 20°C. **(F)** Fluorescence microscopy of the DRE mutant at 37°C. The nucleoid was labelled with Hbs-GFP (green) and the cell membrane was labelled with Nile Red (red).

To identify key nucleotide positions within this ssDNA sequence, thymidine bases were systematically inserted within an inert dC18 substrate and DnaD binding was assessed using fluorescence polarization. The results indicate that two motifs of 5’-TnnT-3’ are necessary and sufficient for DnaD to associate specifically with the ssDNA substrate (Fig. 6C, S15C-D). The symmetry of these repeated motifs suggest that DnaD may bind to ssDNA as a dimer, consistent with the observation that monomeric DnaD^L22A^ cannot bind ssDNA (Fig. 5G). Based on these properties, we have termed the ssDNA sequence complementary to the DnaA-trios the *D*naD *R*ecognition *E*lement (DRE), and we propose that the 5’-TnnT-3’ motifs are critical for DnaD binding.

### Disrupting the DRE impairs cell viability

The DnaA-trios and the DRE appear to be inherently linked ssDNA binding motifs, in which each could potentially be recognized by a distinct replication initiation protein, DnaA and DnaD, respectively. Previous studies have suggested that the DnaA-trios closest to the DnaA-boxes are critical for DnaA unwinding activity (Jaworski et al., 2021; Pelliciari et al., 2021; Richardson et al., 2016; Richardson et al., 2019). Therefore, we hypothesized that mutating the 5’-TnnT-3’ motif furthest from the DnaA-boxes might preferentially inhibit DnaD activity while leaving DnaA relatively unperturbed. Consistent with this notion, using an *in vitro* DnaA strand separation assay it was observed that DnaA activity was not compromised when the distal 5’-TnnT-3’ motif was changed to 5’-AnnA-3’ (Fig. S15E, note that these mutations alter the two distal DnaA-trios).

Guided by the *in vitro* results, a strain was engineered to mutate the 5’-TnnT-3’ motif furthest from the DnaA-boxes within the DRE (5’-CTACTATTACTTC<u>T</u>AC<u>T</u>A-3’ → 5’-CTACTATTACTTC<u>A</u>AC<u>A</u>A-3’). We observed that this mutant displays a growth defect at 20**°**C (Fig. 6D). Fluorescence microscopy showed that the DRE mutant contains fewer *oriC* per nucleoid (Fig. 6E and S15F) and an increase in the number of cells lacking DNA (Fig. 6F and S15G). Marker frequency analysis confirmed that the DRE mutant has a lower DNA replication initiation frequency compared to wild-type cells (Fig. S15H). Taken together, the results are consistent with the DRE functioning as a ssDNA binding site within the *B. subtilis* chromosome origin unwinding region.

## DISCUSSION

The molecular basis for how bacteria recruit a pair of helicases at their chromosome origin to promote bidirectional DNA replication is unknown. Characterization of the essential DNA replication initiation protein DnaD (summarized in Fig. 7A) identified a new ssDNA binding motif (DRE) within the *B. subtilis* chromosome origin. The location of the DRE suggests a mechanism for directing helicase loading to support bidirectional DNA replication.

**Figure 7.**
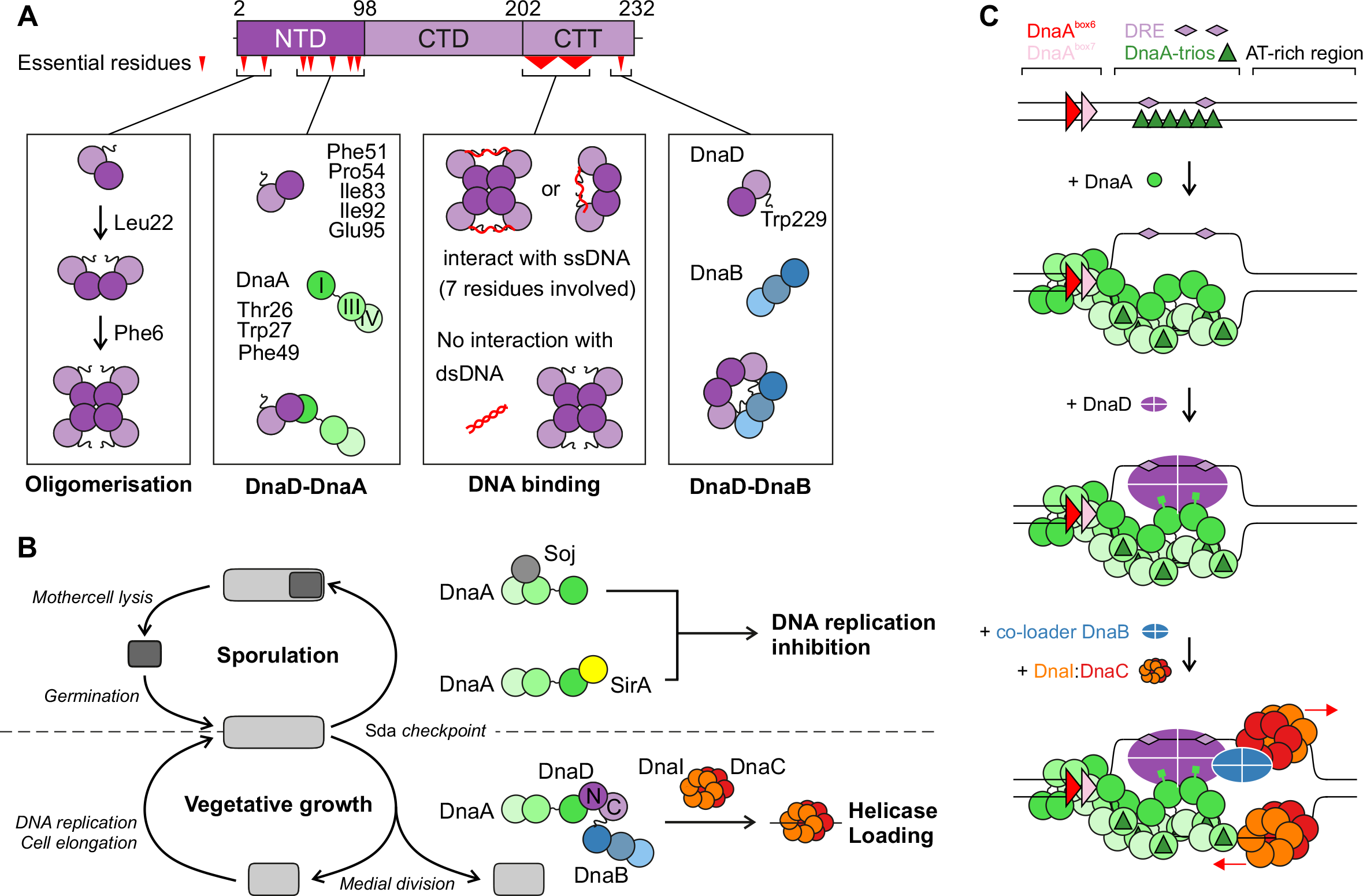
DnaD activities and interactions during DNA replication initiation at *oriC* culminate with it binding the DRE. **(A)** The DnaD functional analysis identified key activities that regulate protein-protein and protein-DNA interactions. **(B)** Regulation of helicase loading by SirA during *B. subtilis* spore development. **(C)** Model of chromosomal replication initiation in *B. subtilis*. DnaA binds DnaA-boxes in the *incC* region, leading to oligomer formation on DnaA-trios and DNA strand separation. DnaD is then loaded on the exposed top strand and provides a pathway for strand-specific helicase recruitment to promote bidirectional DNA replication.

### A mechanism for bidirectional DNA replication at a bacterial chromosome origin

Based on structural and biochemical studies, a model was proposed for helicase loading at one end of a DnaA oligomer, where the AAA+ class of helicase chaperone (DnaI in *B. subtilis,* DnaC in *E. coli*) engages the AAA+ motif of DnaA and guides deposition of a helicase onto ssDNA (Fig. 7B) (Mott et al., 2008). This mechanism results in helicase loading around ssDNA in the correct orientation for 5′◊3′ translocation.

In contrast, the mechanism for loading a helicase onto the complementary strand with the correct geometry was unclear, although it was suggested that the interaction between DnaA^DI^ and helicase^NTD^ might be sufficient. In *E. coli*, biochemical and genetic assays have suggested that DnaA domain I interacts directly with the helicase (Seitz et al., 2000; Sutton et al., 1998). Moreover, it has been shown that DnaA^E21^ is essential for *E. coli* viability and required for helicase loading *in vitro* (Abe et al., 2007). Here in *B. subtilis* we identified residues in DnaA^DI^ that are essential for cell viability and required for the direct recruitment of DnaD to *oriC*. Generalizing, it appears that DnaA^DI^ is critical for helicase recruitment in diverse bacterial species, albeit through different mechanisms. Importantly however, these protein:protein interactions alone do not resolve how a second helicase is recruited to a specific DNA strand for bidirectional replication.

Many bacterial chromosome origins encode a core set of sequence elements that direct DnaA oligomerization onto ssDNA, thus dictating the strand onto which the AAA+ helicase chaperone docks (Pelliciari et al., 2021). Here we report that the DRE, located opposite to where the DnaA oligomer binds, provides a mechanism for orchestrating strand-specific DnaD recruitment. Considered together with studies of primosome assembly at a single-strand origin *in vivo* where binding of DnaD to ssDNA promoted subsequent helicase loading (Bruand et al., 1995) (Fig. S16), we propose that the specific interaction of DnaD with the DRE provides a pathway for loading a second helicase to support bidirectional DNA replication.

The proposed model for DnaD recruitment raises several fundamental questions. How does DnaD (along with DnaA and DnaB) orientate the DnaI:helicase complex? We speculate that within the open complex formed at *oriC*, the junction between dsDNA and ssDNA could direct the subsequent events. Additionally, the nucleoprotein complexes formed at *oriC* between DnaA, DnaD and DnaB could play a role.

How is the temporal loading of two helicases orchestrated? Studies of helicase loading onto artificial DNA scaffolds that mimic an open origin indicated that DnaA preferentially recruits helicase onto the strand corresponding to where the DRE is located (Weigel and Seitz, 2002). Whether this order of recruitment holds during the physiological helicase loading reaction is unclear. While we favour a model where loading of the two helicases at *oriC* is reproducibly sequential, an alternative hypothesis is that loading of the two helicases is stochastic.

Are there other sequence elements within *oriC* that direct helicase loading? The discovery of the DnaA-trios and the DRE indicate that bacterial chromosome origins encode more information than previously appreciated. We note that many chromosome origins contain an intrinsically unstable AT-rich region (Kowalski and Eddy, 1989; Krause et al., 1997) where one of the helicases is loaded *in vitro* (Fang et al., 1999). We wonder whether additional sequence dependent information may be located within these AT-rich sites, or elsewhere. Further characterization of the nucleoprotein complexes formed at *oriC*, as well as dissection of downstream helicase loader proteins, will be needed to provide answers.

### Regulation of helicase recruitment in bacteria

Here we report that the essential DnaA^DI^-DnaD^NTD^ interface is targeted by the inhibitor of DNA replication initiation SirA, which binds DnaA^DI^ and occludes DnaD (Fig. 4E). To our knowledge, this is the first description of a bacterial mechanism for regulating helicase recruitment. Interestingly, while the interaction of SirA with DnaA^DI^ inhibits helicase recruitment at the chromosome origin during endospore development (Fig. 7C), this regulatory system would not perturb the interaction of DnaD with the replication restart primosome (Huang et al., 2016) required to ensure completion of genome replication (Fig. S16).

While SirA is the first example of an endogenous bacterial system that regulates helicase recruitment, we note that diverse viruses effectively hijack the bacterial helicase loading pathway during their infective life cycle (Hood and Berger, 2016; Kimura et al., 2010; Klein et al., 1980; Noguchi and Katayama, 2016; Odegrip et al., 2000). Homologs of *dnaD* are present in the majority of *Firmicutes* including several clinically relevant human pathogens such as *Staphylococcus, Streptococcus, Enterococcus,* and *Listeria* (Briggs et al., 2012). Moreover, replication of *S. aureus* multiresistant plasmids have been shown to require an initiation protein with structural homology to DnaD (Schumacher et al., 2014). Therefore, the multiple essential activities of DnaD, combined with the appreciation that helicase recruitment and loading mechanisms in bacteria and eukaryotes are distinct, indicates that DnaD homologs are an attractive target for antibacterial drug development.

## METHODS

### Media and chemicals

Nutrient agar (NA; Oxoid) was used for routine selection and maintenance of both *B. subtilis* and *E. coli* strains. For experiments in *B. subtilis* cells were grown using Luria-Bertani medium (LB). Supplements were added as required: ampicillin (100 µg/ml), chloramphenicol (5 µg/ml), kanamycin (5 µg/ml), spectinomycin (50 µg/ml). All chemicals and reagents were obtained from Sigma-Aldrich unless otherwise noted.

### Microscopy

To visualize cells during the exponential growth phase, starter cultures were grown in imaging medium (Spizizen minimal medium supplemented with 0.001 mg/mL ferric ammonium citrate, 6 mM magnesium sulphate, 0.1 mM calcium chloride, 0.13 mM manganese sulphate, 0.1% glutamate, 0.02 mg/mL tryptophan) with 0.5% glycerol, 0.2% casein hydrolysate and 0.1 mM IPTG. Saturated cultures were diluted 1:100 into fresh imaging medium supplemented with 0.5% glycerol and 0.1 mM IPTG and allowed to grow for three mass doublings. Early log cells were then spun down for 5 minutes at 9000 rpm, resuspended in the same medium lacking IPTG and further incubated for 90 minutes before imaging.

Cells were mounted on ∼1.4% agar pads (in sterile ultrapure water) and a 0.13- to 0.17-mm glass coverslip (VWR) was placed on top. Microscopy was performed on an inverted epifluorescence microscope (Nikon Ti) fitted with a Plan Apochromat Objective (Nikon DM 100x/1.40 Oil Ph3). Light was transmitted from a CoolLED pE-300 lamp through a liquid light guide (Sutter Instruments), and images were collected using a Prime CMOS camera (Photometrics). The fluorescence filter sets were from Chroma: GFP (49002, EX470/40 (EM), DM495lpxr (BS), EM525/50 (EM)) and mCherry (49008, EX560/40 (EM), DM585lprx (BS), EM630/75 (EM)). Digital images were acquired using METAMORPH software (version 7.7) and analysed using Fiji software (Schindelin et al., 2012). All experiments were independently performed at least twice, and representative data are shown.

The number of origins was quantified using the Trackmate plugin within the Fiji software (Tinevez et al., 2017). Background was subtracted from fluorescence images set to detect 8-10 pixel blob diameter foci over an intensity threshold of 150 relative fluorescence units. A mask containing the detected origin foci was created and merged with the nucleoids channel, and the number of origins per nucleoid was determined and averaged for a minimum of 100 cells from each strain that was examined. For origin and *dnaD^7A^* mutants, the count of nucleoids was determined using line plots of a 10 pixel width drawn across the length of cells. For each field of view, an average of the whole cell fluorescence was measured and used to normalise individual line plots, and the exact number of nucleoids per cell was assessed by counting the number of peaks crossing the zero line.

### Phenotype analysis of dnaA mutants using the inducible oriN strain

Strains were grown for 48 hours at 37°C on NA plates supplemented with kanamycin either with or without IPTG (0.1 mM). All experiments were independently performed at least twice and representative data is shown.

### Phenotype analysis of dnaD mutants using the inducible dnaD-ssrA strain

Strains were grown for 18 hours at 37°C on NA plates (spot-titre assays) or in Penassay Broth (PAB, plate reader experiments) either with or without IPTG (0.1 mM). All experiments were independently performed at least twice and representative data is shown.

### Phenotype analysis of origin mutants

Strains were grown for 72 hours at 20°C or 37°C on NA plates. All experiments were independently performed at least twice and representative data is shown.

### Bacterial two-hybrid assay

*Escherichia coli* strain HM1784 was transformed using a combination of complementary plasmids and grown to an OD600nm of 0.5 in LB containing ampicillin and spectinomycin, before diluting 1:10,000 and spotting onto nutrient agar plates containing antibiotics and the indicator X-gal (0.008%). Plates were incubated at 30°C for 48 hours and imaged using a digital camera. Experiments were independently performed at least twice and representative data is shown.

### Immunoblot analysis

Proteins were separated by electrophoresis using a NuPAGE 4-12% Bis-Tris gradient gel run in MES buffer (Life Technologies) and transferred to a Hybond-P PVDF membrane (GE Healthcare) using a semi-dry apparatus (Bio-rad Trans-Blot Turbo). DnaA, DnaD and FtsZ were probed with polyclonal primary antibodies (Eurogentec) and then detected with an anti-rabbit horseradish peroxidase-linked secondary antibody using an ImageQuant LAS 4000 mini digital imaging system (GE Healthcare). Detection of DnaA, DnaD and FtsZ was within a linear range. Experiments were independently performed at least twice and representative data is shown.

### Pull-down assay of His6-DnaA^DI^-DnaD^NTD^ complexes

BL21 (DE3) *E. coli* cells containing the different expression plasmids (pSP075, pSP080, pSP081, pSP082, pSP083 and pSP085) were grown overnight in 5 ml of LB supplemented with kanamycin at 37°C. The following day cells were diluted in 50 ml of fresh medium until A600 reached 0.5. Protein expression was induced by adding 1 mM IPTG for 4 hours at 30°C. Cells were collected by centrifugation and resuspended in 2 ml of resuspension buffer (30 mM Hepes pH 7.5, 250 mM potassium glutamate, 10 mM magnesium acetate, 20% sucrose, 30 mM imidazole) supplemented with 1 EDTA-free protease inhibitor tablet. Bacteria were lysed with two sonication cycles at 10 W for 3 minutes with 2 second pulses. Cell debris were pelleted by centrifugation at 25,000 g at 4°C for 30 minutes and the supernatant was filtered through 0.2 µm filters. The clarified lysate was then loaded onto Ni-NTA spin columns (QIAgen) and proteins purified according to manufacturer protocol washing the column with Washing Buffer (30 mM Hepes pH 7.5, 250 mM potassium glutamate, 10 mM magnesium acetate, 20% sucrose, 100 mM imidazole) and eluting bound proteins with elution buffer (30 mM Hepes pH 7.5, 250 mM potassium glutamate, 10 mM magnesium acetate, 20% sucrose, 1 M imidazole). The eluates were loaded on a NuPAGE 4-12% Bis-Tris gradient gel run in MES buffer (Life Technologies) and analysed with IstantBlue staining (Merck).

### ChIP-qPCR

Chromatin immunoprecipitation and quantitative PCR were performed as previously described (Fisher et al., 2017) with minor modification (see Supplementary Methods).

### Cryo-EM Sample Preparation and Data Collection

Four-microliter samples of purified wild-type DnaD were applied to plasma-cleaned Ultrafoil 2/2 200 grids, followed by plunge-freezing in liquid ethane using a Leica EM GP. Data collection was carried out at liquid nitrogen temperature on a Titan Krios microscope (Thermo Fisher Scientific) operated at an accelerating voltage of 300 kV. Micrograph movies were collected using EPU software (FEI) on a Gatan K3 detector in counting mode with a pixel size of 0.67 Å. A total of 3095 movie frames were acquired with a defocus range of approximately -0.7 to -2.7 μm. Each movie consisted of a movie stack of 30 frames with a total dose of 50 electron/Å^2^ over 1.5 seconds with a total dose of ∼50 electron/Å^2^ at a dose rate of 15 electron/pixel/second.

### Cryo-EM Image Processing, reconstruction and model fitting

The movie stacks were aligned and summed with dose-weighting using MotionCor2 (Zheng et al., 2017). Contrast transfer function (CTF) was estimated by CtfFind4 (Rohou and Grigorieff, 2015), and images with poor CTF estimation were eliminated. A small subset of 200 micrographs were used to pick the particles using the blob picker tool in CryoSparc v3.1.0 (Punjani et al., 2017). These particles were 2D classified to generate a template which was subsequently used for particles picking using the template picket tool in CryoSparc. A total of 1299061 initial particles were picked and subjected to several iterative rounds of 2D classification, removing particles of poor template classes after each round of classification. A final set of 73190 good particles were used to generate an *ab-initio* model using a C2 symmetry, as a clear two-fold symmetry was visible from the 2D classes (Fig 2E). Using the *ab-initio* model, the particles were subjected to 3D homogenous refinement in CryoSparc which yielded a map at 10.1 Å at 0.5 FSC cut off.

A poly alanine model of the crystal structures of the DnaD^NTD^ (PDB 2V79) and the DnaD^CTD^ (PDB 2ZC2) were used to dock into the cryo-EM map of DnaD. The C-terminal domain can be readily identified and was docked into the density using Chimera (Pettersen et al., 2004). The entire N-terminal domain could not be readily docked in to the cryo-EM density hence a flexible fitting approach was adopted. The β-hairpin density could be easily identified within the map which was docked in the density first. The remainder of the structure was manually fitted in the density using Coot (Emsley and Cowtan, 2004). A single round of real space refinement was performed to remove any clashes and idealize the model using Phenix (Liebschner et al., 2019).

## ACKNOWLEDGEMENTS

SEC-MALLS experiments were performed by Dr. Andrew Leech at the Molecular Interaction laboratory as a service from the University of York. Research support was provided to HM by a Wellcome Trust Senior Research Fellowship (204985/Z/16/Z) and a grant from the Biotechnology and Biological Sciences Research Council (BB/P018432/1). Research support was provided to PS by a grant from the Biotechnology and Biological Sciences Research Council (BB/R013357/1). DS was supported by a Research Excellence Academy Studentship from the Faculty of Medical Sciences at Newcastle University. EM was supported by Erasmus+.

## AUTHOR CONTRIBUTIONS

CW, DS, SF, SP, EM, NBC, PS, TRDC, AI, HM contributed to the conception/design of the work. CW, DS, SF, SP, NBC, AI, HM generated results presented in the manuscript. NBC collected the cryo-EM data, TRDC and AI processed the cryo-EM data. CW, DS, SF, SP, AI created Figures. HM, CW, AI wrote the manuscript. HM, CW, DS, SF, SP, PS, AI edited the manuscript.

---

**Figure S1.**
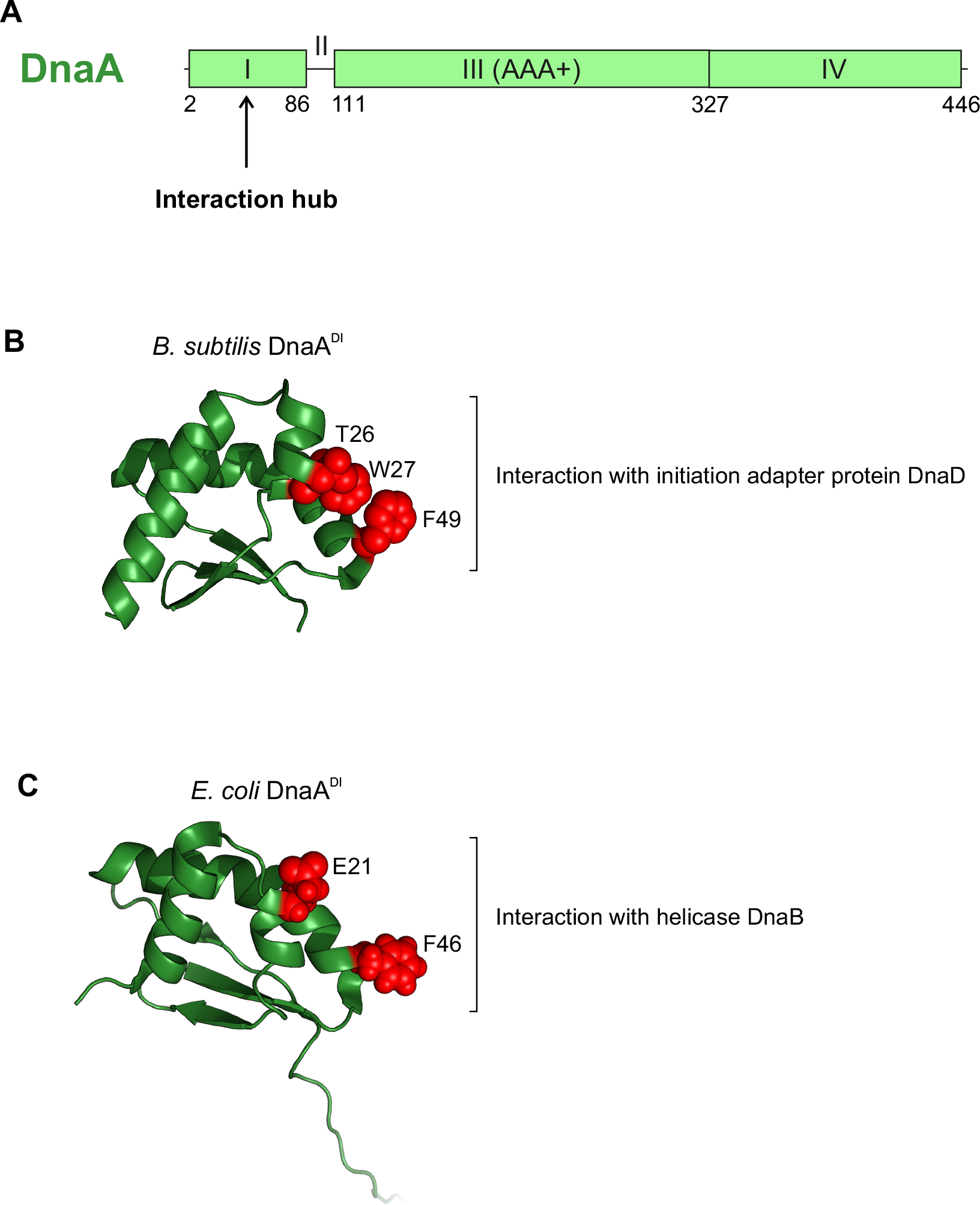
Domain organisation of DnaA highlighting a shared interaction hub in domain I. **(A)** *B. subtilis* Dna A domain organisation with amino acid boundaries indicated. **(B)** Crystal structure of *B. subtilis* DnaA domain I (PDB 4TPS) with residues thought to be involved in protein-protein interactinos highlighted in red. **(C)** NMR structure of *E. coli* DnaA domain I (PDB 2E0G) with residues thought to be involved in protein-protein interactions highlighted in red.

**Figure S2.**
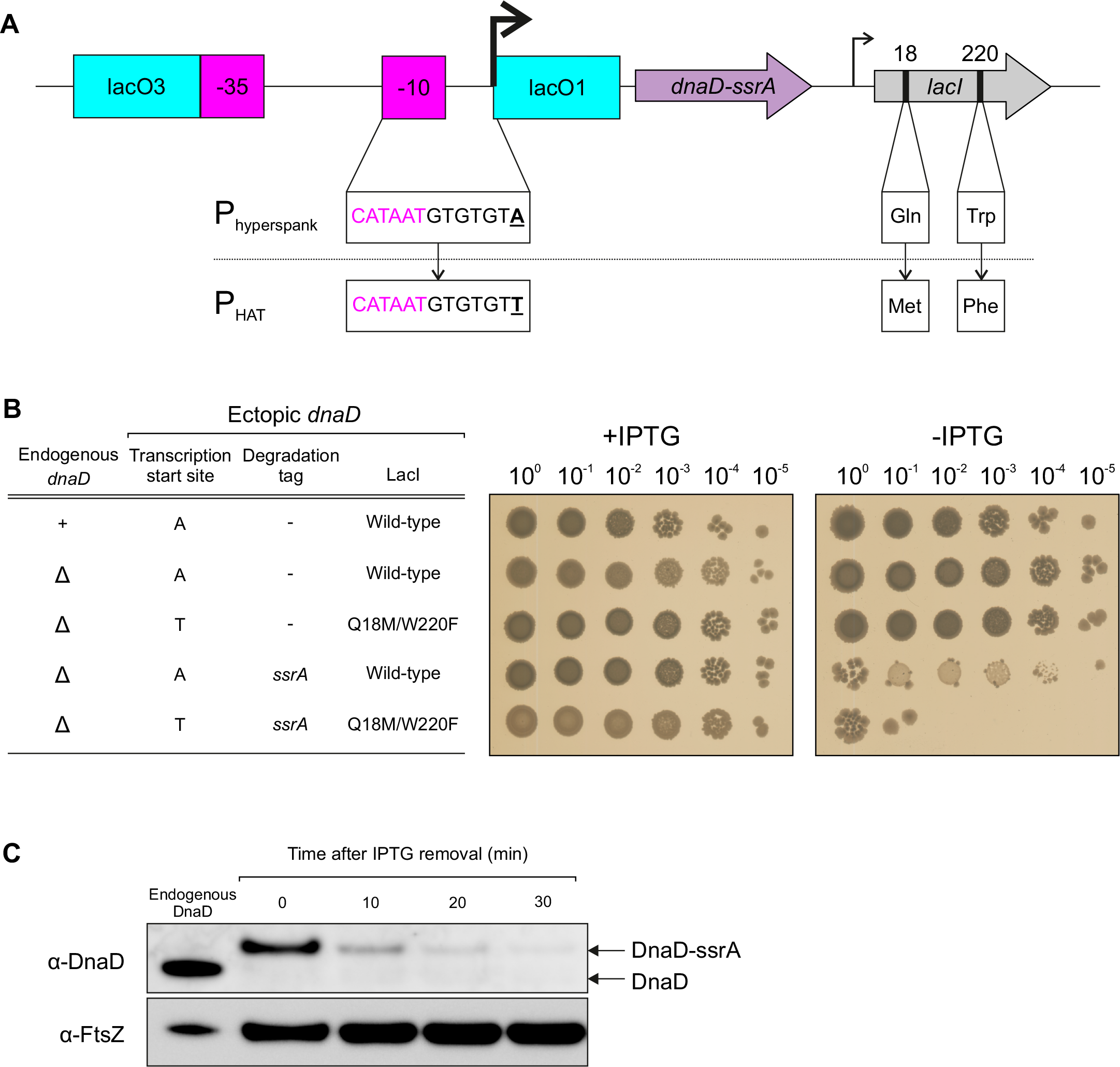
Construction of the inducible *dnaD-ssrA* strain. **(A)** Schematics of the inducible system used to drive the expression of the *dnaD-ssrA* fusion. **(B)** Spot-titre assay showing the combination required to achieve conditional DnaD-ssrA complementation. *dnaD P_HYPERSPANK_-dnaD* (CW2), *ΔdnaD P_HYPERSPANK_-dnaD* (CW231), *ΔdnaD P_HAT_-dnaD* (CW103), *ΔdnaD P_HYPERSPANK_-dnaD-ssrA* (CW232), *ΔdnaD P_HAT_-dnaD-ssrA* (CW164). **(C)** Immunoblot analysis of the inducible *dnaD-ssrA* cassette (CW197); endogenous *dnaD* control (HM715). The tubulin homolog FtsZ was used as a loading control.

**Figure S3.**
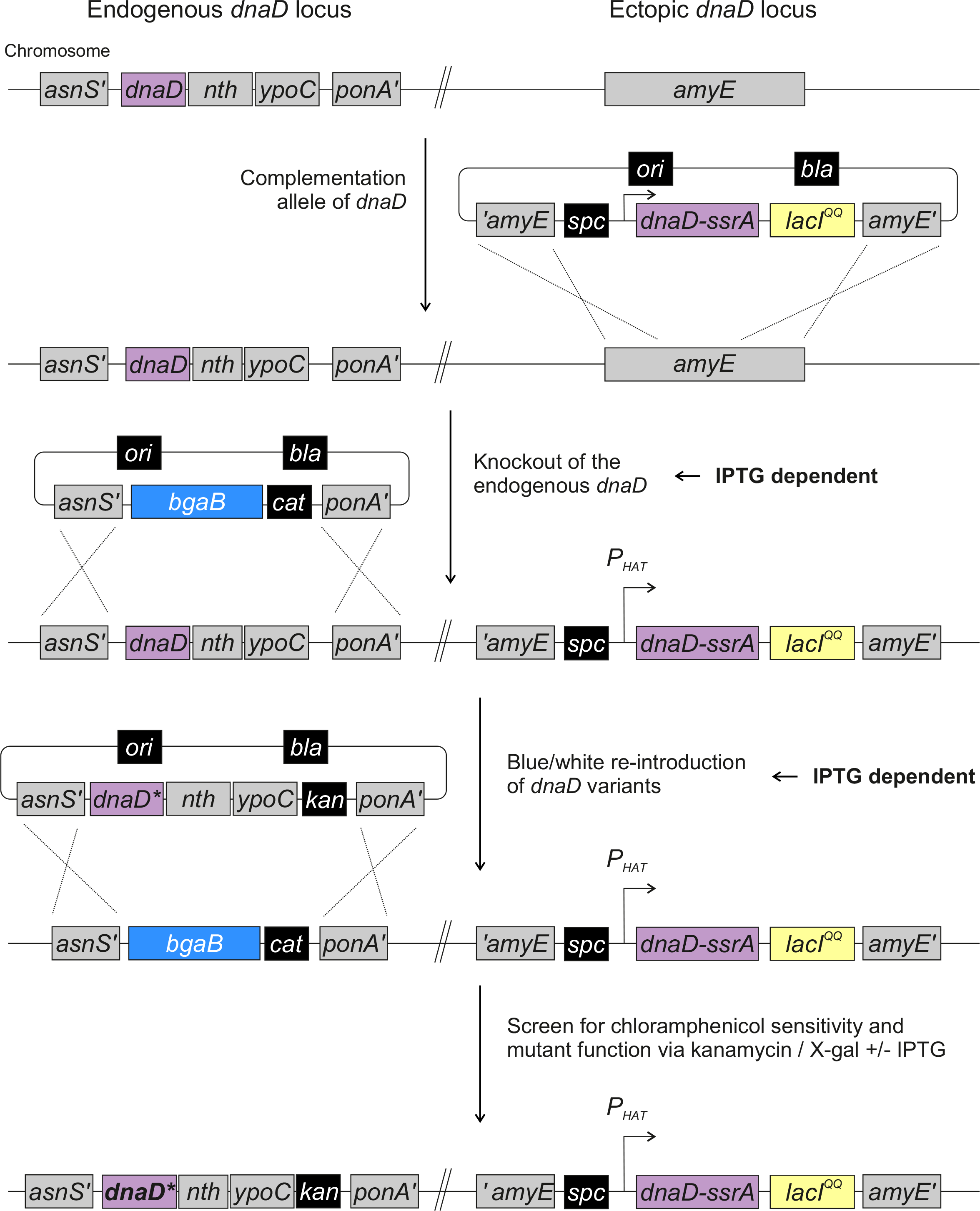
Methodology for genetic complementation and introduction of *dnaD* mutants. Schematics of the blue/white screening assay. The inducible complementation cassette *dnaD-ssrA* was inserted at the *amyE* locus, followed by replacement of the native *dnaD* operon by a *bgaB* cassette (encoding β-galactosidase). Selection of *dnaD* mutants is performed in the presence of kanamycin, X-gal (blue/white) and IPTG (functional complementation).

**Figure S4.**
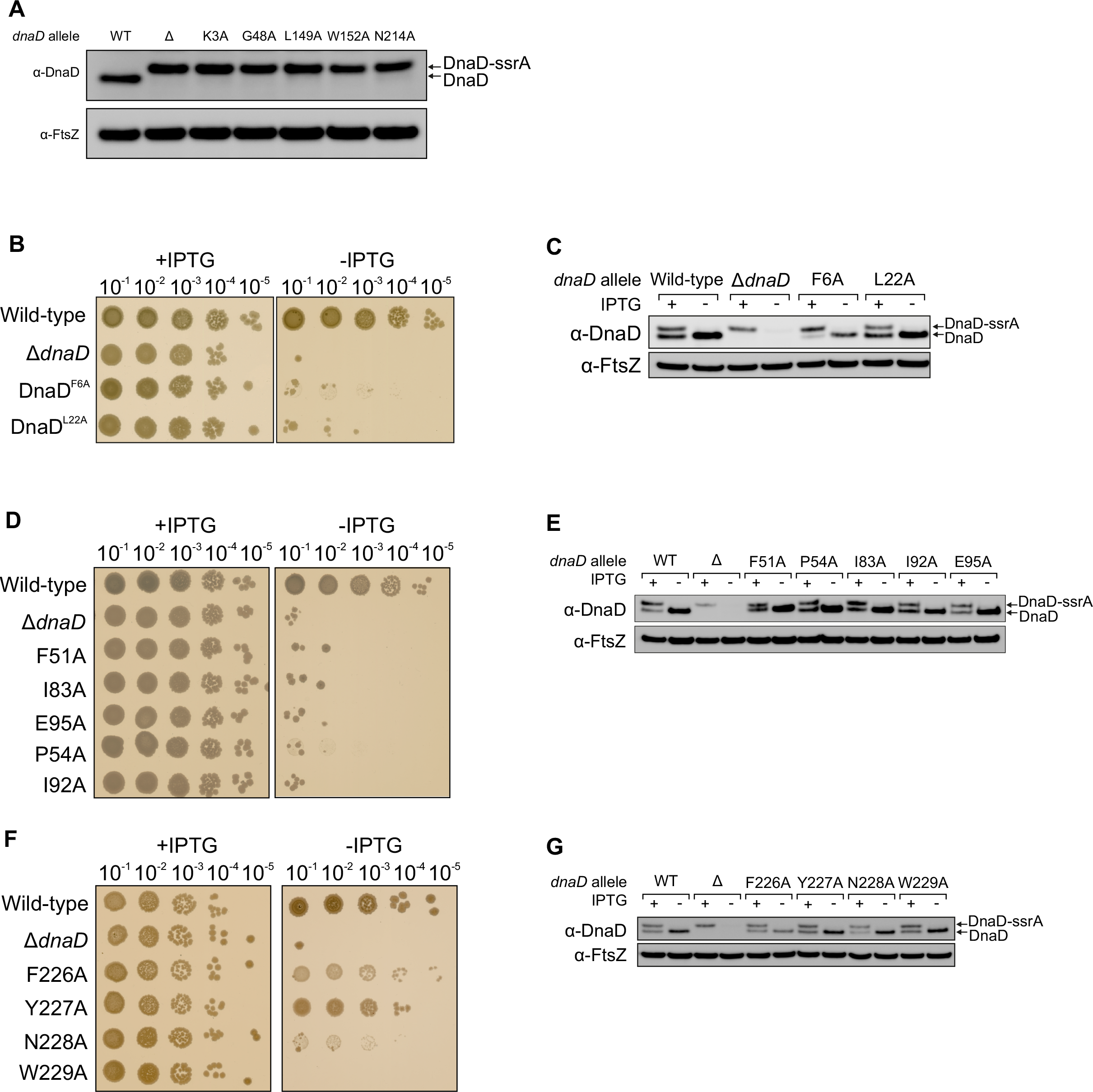
Analysis of lethal and sick alanine substitutions in DnaD. **(A)** Immunobloting shows that some lethal alanine substitutions in DnaD were not well expressed *in vivo*. **(B, D, F)** Spot titre analysis of alanine substitutions in DnaD that were well expressed as judged by immunoblotting **(C, E, G)**. The tubulin homolog FtsZ was used as a loading control.

**Figure S5.**
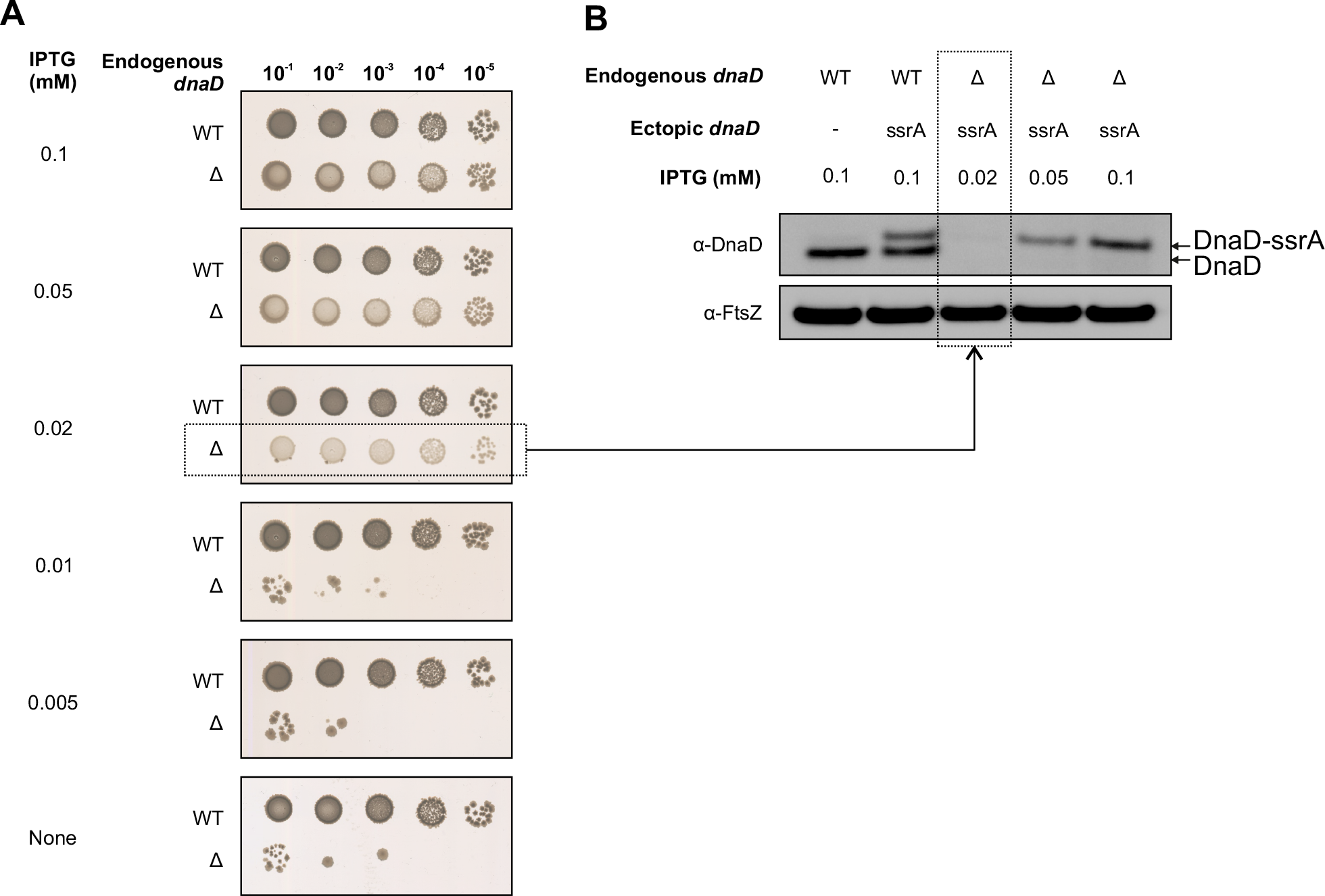
Low levels of DnaD expression sustain cell growth. **(A)** DnaD-SsrA was titrated via IPTG induction. The *dnaD-ssrA* cassette was able to sustain growth at IPTG concentration of 0.02 mM and above. **(B)** Immunobloting shows that expression of DnaD was undetectable in viable colonies grown with 0.02 mM IPTG. The tubulin homolog FtsZ was used as a loading control.

**Figure S6.**
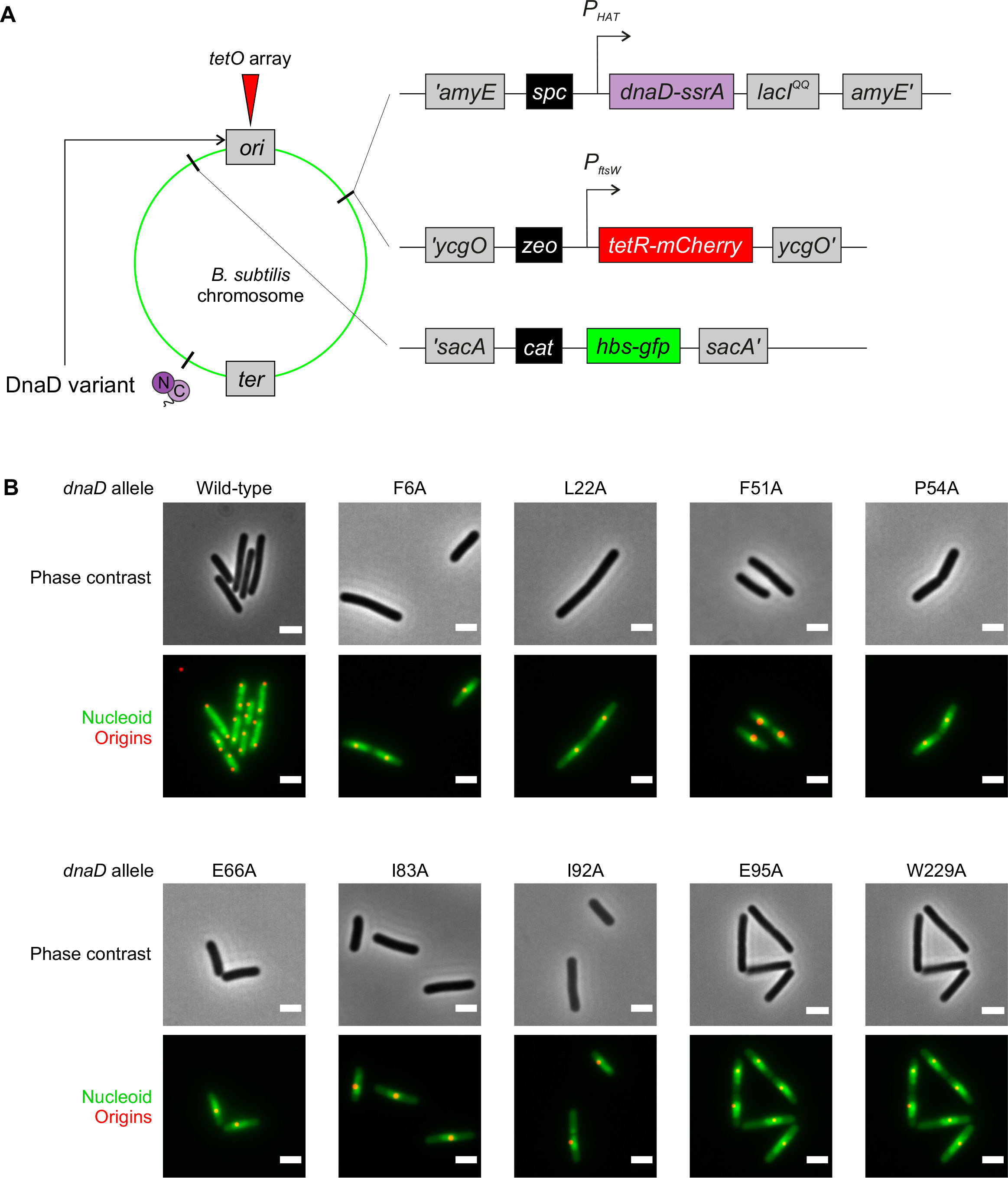
Methodology for single-cell analysis of *dnaD* mutants using fluorescence microscopy. **(A)** Schematics of the dual fluorescence system with TetR-mCherry binding to *tetO* sites located near the origin and Hbs-GFP allowing visualisation of the nucleoid. **(B)** Representative images of essential *dnaD* mutants observed by fluorescence microscopy via the system described in (A).

**Figure S7.**
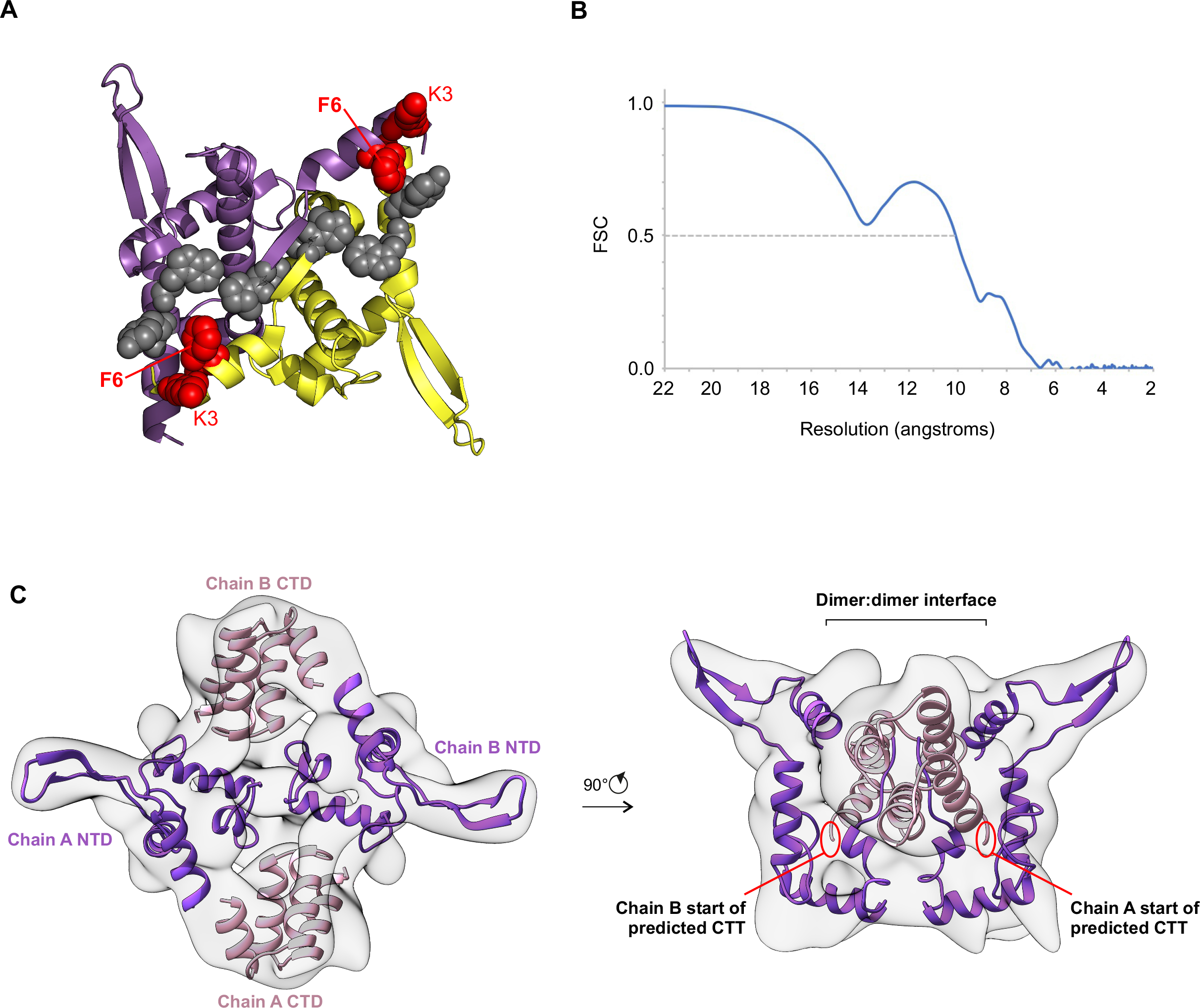
A network of critical residues along the proposed DnaD dimer:dimer interface. **(A)** Crystal structure of DnaD N-terminal domain (PDB 2V79) mapped with alanine substitutions that are either lethal essential (red) or perturb growth (grey). **(B)** Overall resolution of the DnaD dimer, derived from two independently refined half-maps, using the FSC=0.5 criteria. **(C)** Cryo-EM map fitted with the available DnaD structures (purple N-terminal domains from PDB 2V79 and pink C-terminal domains from 2ZC2).

**Figure S8.**
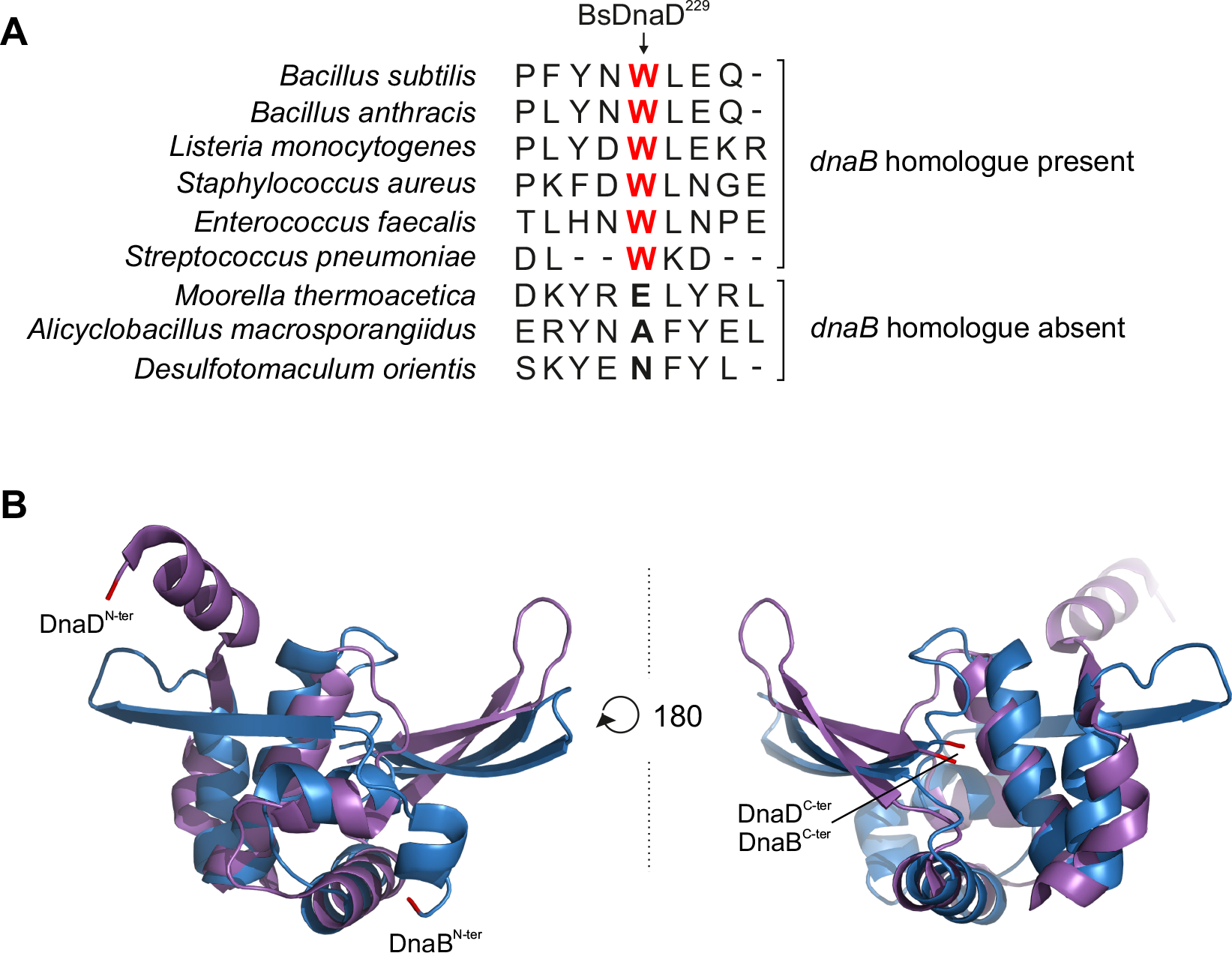
Analysis of the DnaD C-terminal tail. **(A)** *B. subtilis* DnaD^W229^ (BsDnaD^229^) is conserved in homologs that also encode a copy of *dnaB*. **(B)** Structural overlap between DnaD (purple) and DnaB (blue) crystal structures (respectively PDB 2V79 and 5WTN).

**Figure S9.**
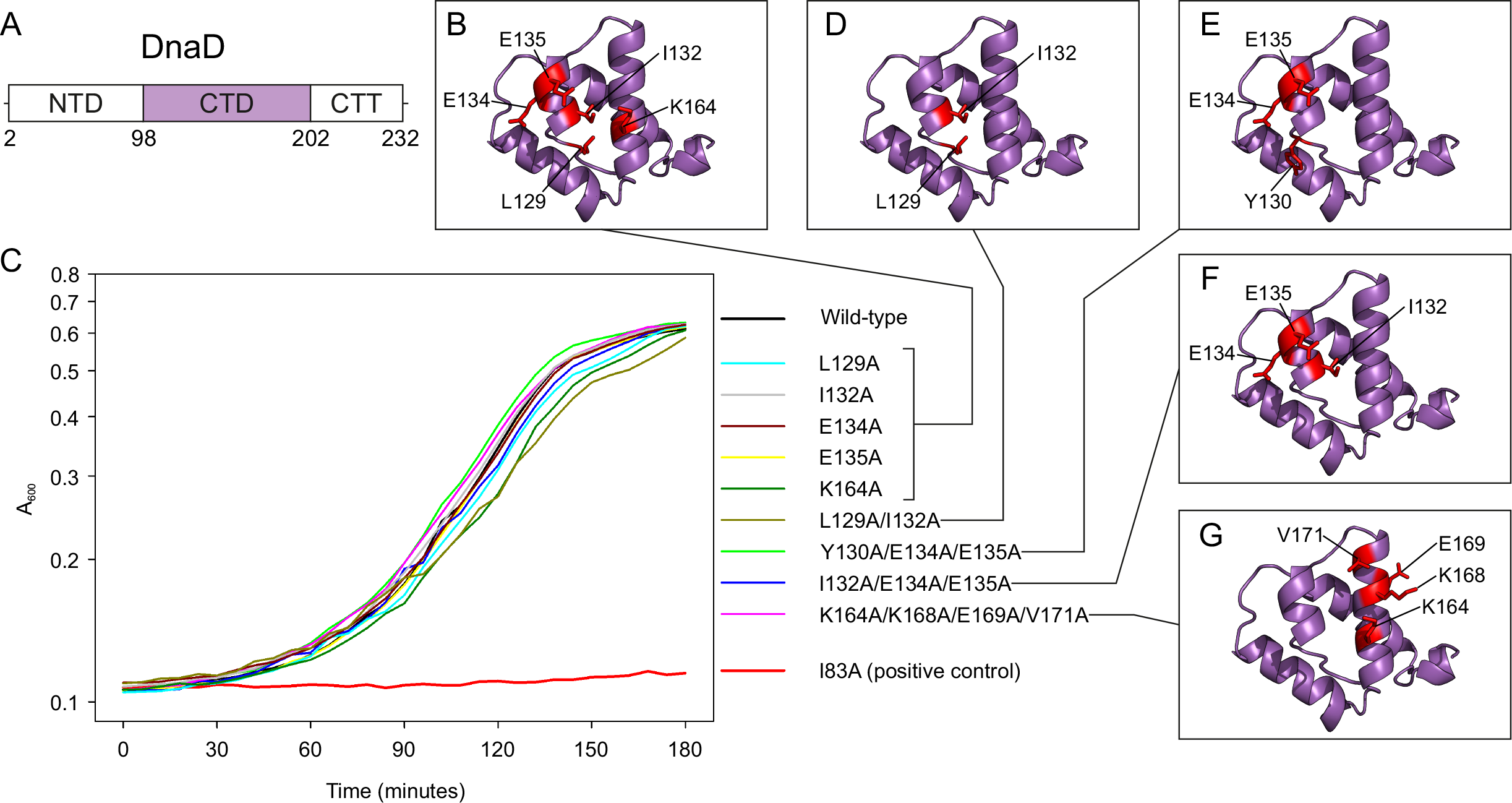
Residues in DnaD C-terminal domain that interact with DnaA^DI^ *in vitro* are not essential *in vivo*. **(A)** Domain organisation of DnaD with amino acid boundaries indicated. **(B)** Individual substitutions in DnaD^CTD^ mapped onto the NMR structure. **(C)** Growth analysis of *B. subtilis* DnaD variants using the inducible *dnaD-ssrA* strain. Wild-type (CW162); *dnaD^L129A^* (CW179), *dnaD^I132A^* (CW167), *dnaD^E134A^* (CW171), *dnaD^E135A^* (CW172), *dnaD^K164A^* (CW173), *_dnaD_L129A/I132A* _(CW176), *dnaD*_*Y130A/E134A/E135A* _(CW177), *dnaD*_*I132A/E134A/E135A* _(CW178),_ *dnaD^K164A/K168A/E169A/V171A^* (CW168) and *dnaD^I83A^* (CW170). **(D-G)** Multiple substitutions in DnaD C-terminal domain mapped onto the NMR structure (Marston et al. 2010).

**Figure S10.**
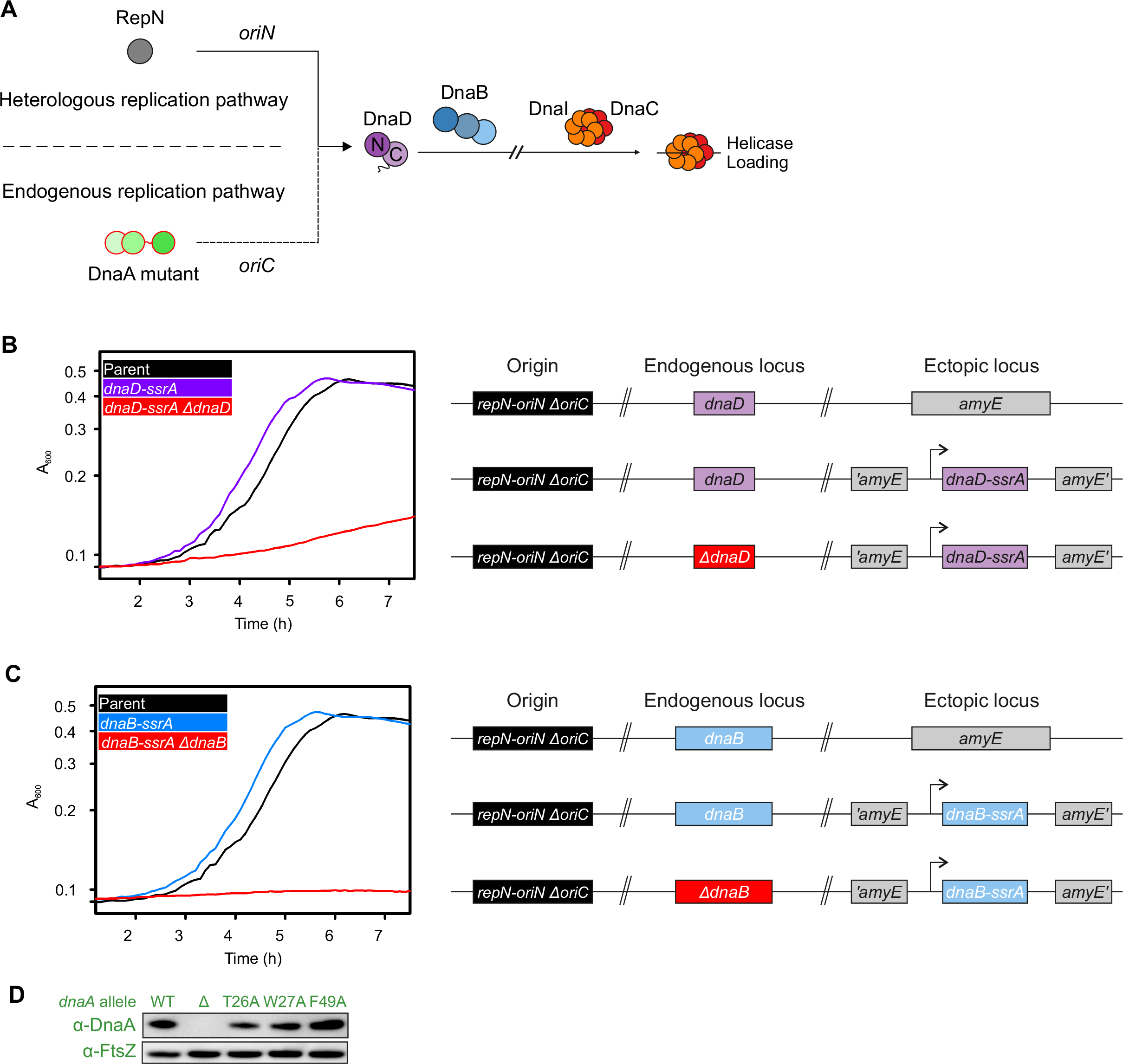
Endogenous and heterologous DNA replication systems for the study of DnaA variants in *B. subtilis*. **(A)** Endogenous replication via DnaA at *oriC* can be complemented by the presence of the heterologous *oriN-repN* replication system. Note that both pathways require DnaD and DnaB to achieve helicase loading. **(B)** Plate reader assay showing growth of a strain replicating exclusively via *oriN,* with and without DnaD expression. **(C)** Plate reader assay showing growth of a strain replicating exclusively via *oriN,* with and without DnaB expression. **(D)** Immunobloting shows that DnaA^DI^ variants were expressed at a similar level to wild-type in the context of the *oriN* strain. The tubulin homolog FtsZ was used as a control.

**Figure S11.**
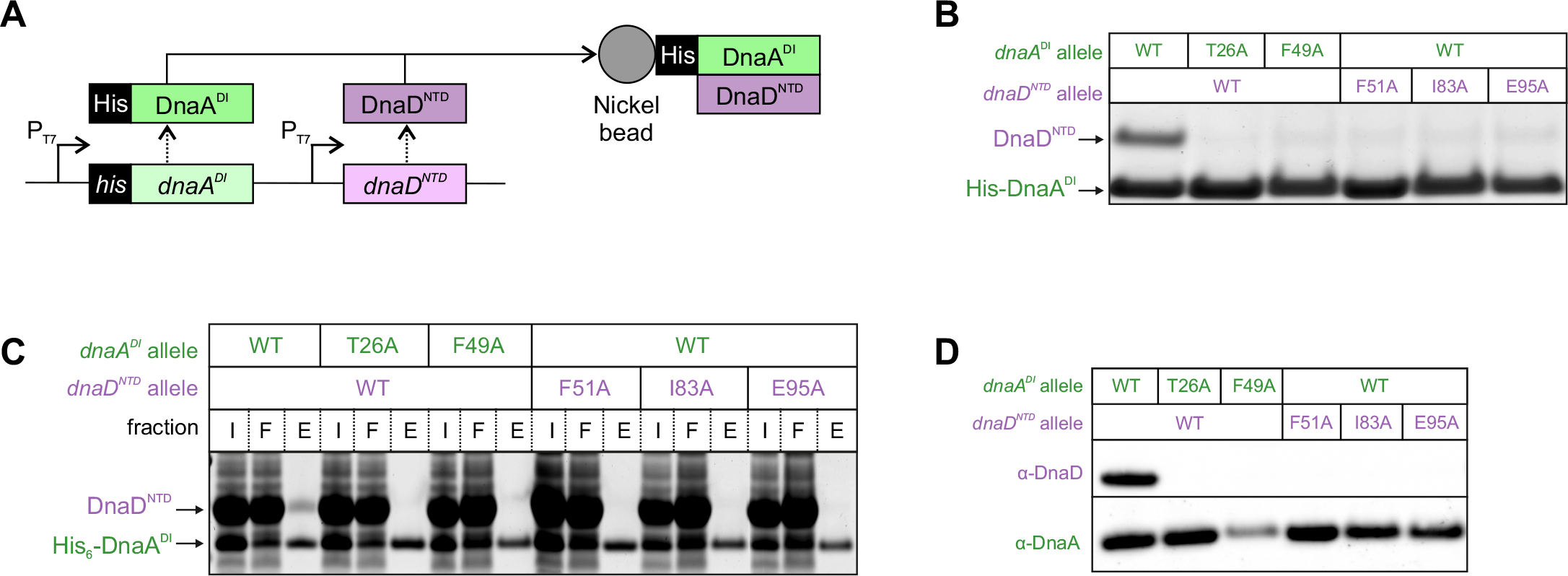
DnaA^DI^ and DnaD^NTD^ mutants disrupt the DnaA-DnaD interaction. **(A)** Schematic of the pull-down assay using *his_6_ -dnaA^DI^* and *dnaD^NTD^.* **(B)** Pull-down assay showing loss of interaction between wild-type and variants of His_6_ -DnaA^DI^ and DnaD^NTD^. Wild type *his*_6_ *-dnaA^DI^/dnaD^NTD^* (pSP75), *his*_6_ *-dnaA^DI-T26A^/dnaD^NTD^* (pSP83), *his*_6_ *-dnaA^DI-F49A^/dnaD^NTD^* (pSP85), *his*_6_ *-dnaA^DI^/dnaD^NTD-F51A^* (pSP80), *his*_6_*-dnaA^DI^/dnaD^NTD-I83A^* (pSP81), *his*_6_*-dnaA^DI^/dnaD^NTD-E95A^* (pSP82). **(C)** Eluate staining from DnaA-DnaD pull-down assays showing loss of interaction between His_6_-DnaA^DI^ and DnaD^NTD^ when using mutant variants. Strains are same as panel (B). Input, Flow through and Eluate fractions are respectively indicated as I, F and E. **(D)** Immunoblot analysis of DnaA and DnaD mutant overexpression eluates from pull-down assays. Strains are same as panel (B).

**Figure S12.**
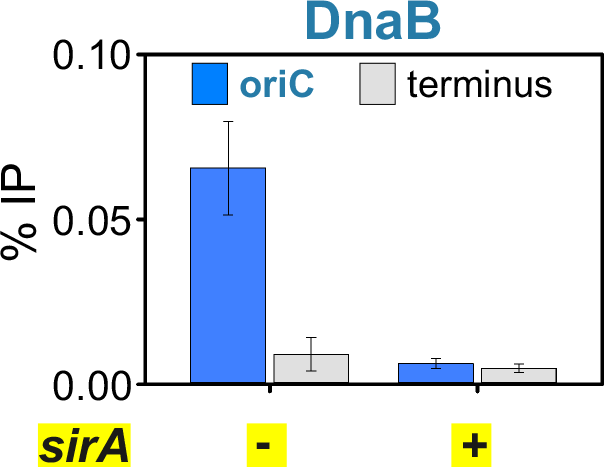
SirA overexpression abolishes DnaB recruitment to *oriC*. ChIP of DnaB at *oriC* following overexpression of SirA.

**Figure S13.**
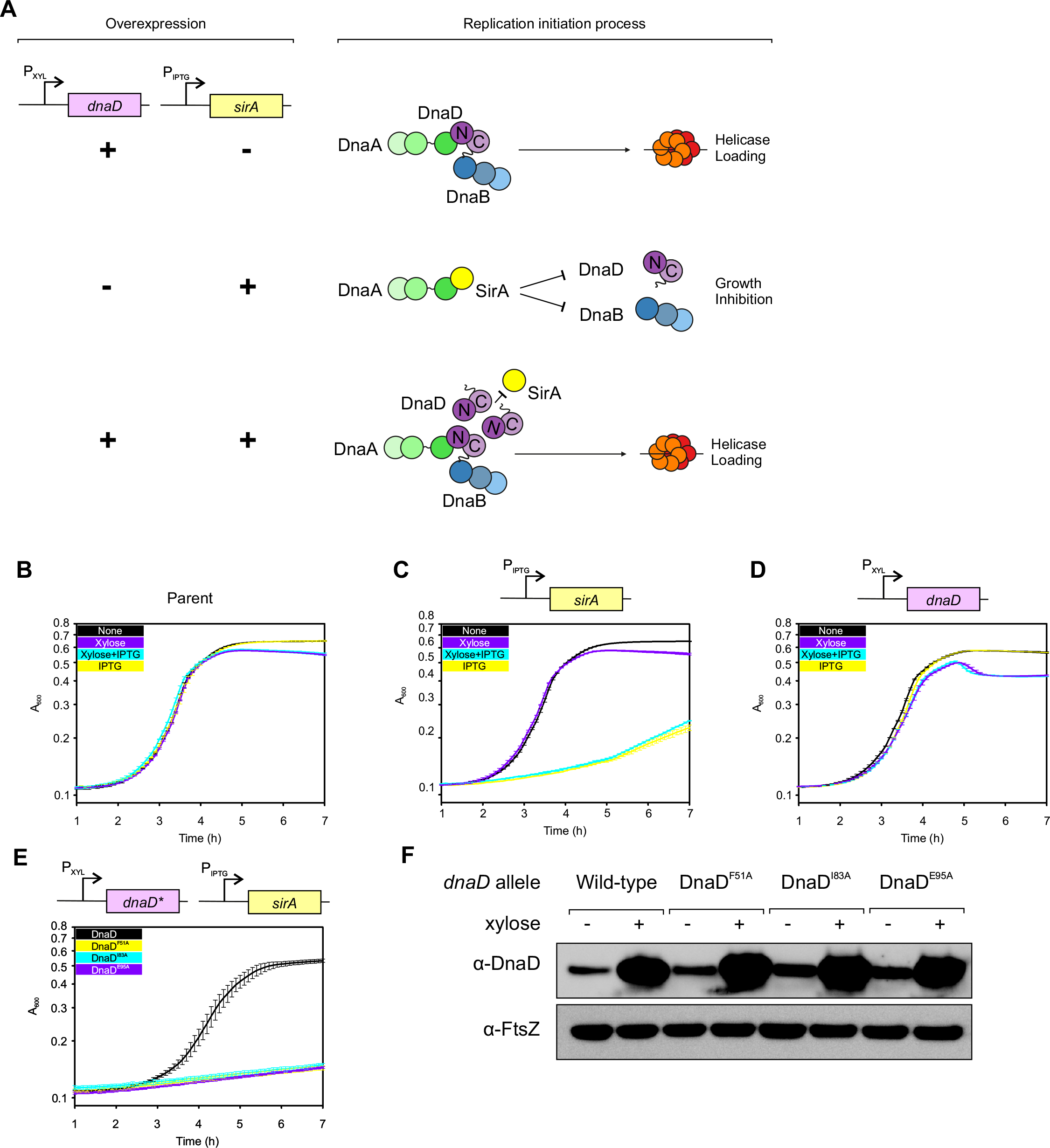
SirA inhibits the DnaA:DnaD interaction by preventing DnaD recruitment to *oriC*. **(A)** Schematics of DnaD/SirA overexpression assay. DnaD overexpression on its own does not affect bacterial growth while SirA overexpression inhibits growth. This inhibition is alleviated when overexpressing DnaD along SirA. **(B-D)** Plate reader analysis with either no inducer (None), Xylose (0.35%), IPTG (0.035 mM) or Xylose (0.35%) and IPTG (0.035 mM). **(B)** Shows that wild type *B. subtilis* (HM715) grows in all conditions. **(C)** Shows that SirA overexpression in a strain background lacking the DnaD overexpression cassette inhibits bacterial growth, and that this inhibition is solely due to the addition of IPTG. *P_HYPERSPANK_-sirA* (CW260). **(D)** Shows that DnaD overexpression in a strain background lacking the SirA overexpression cassette does not affect bacterial growth. *P_XYL_-dnaD* (CW261). **(E)** Plate reader analysis in the presence of Xylose (0.35%) and IPTG (0.1mM) shows that DnaD^NTD^ mutants F51A, I83A and E95A do not rescue SirA-dependent growth inhibition. Error bars in (B-E) indicate the standard error of the mean for two biological replicates. **(F)** Immunoblot analysis of DnaD variants overexpression by xylose induction (0.35%). The tubulin homolog FtsZ was used as a control.

**Figure S14.**
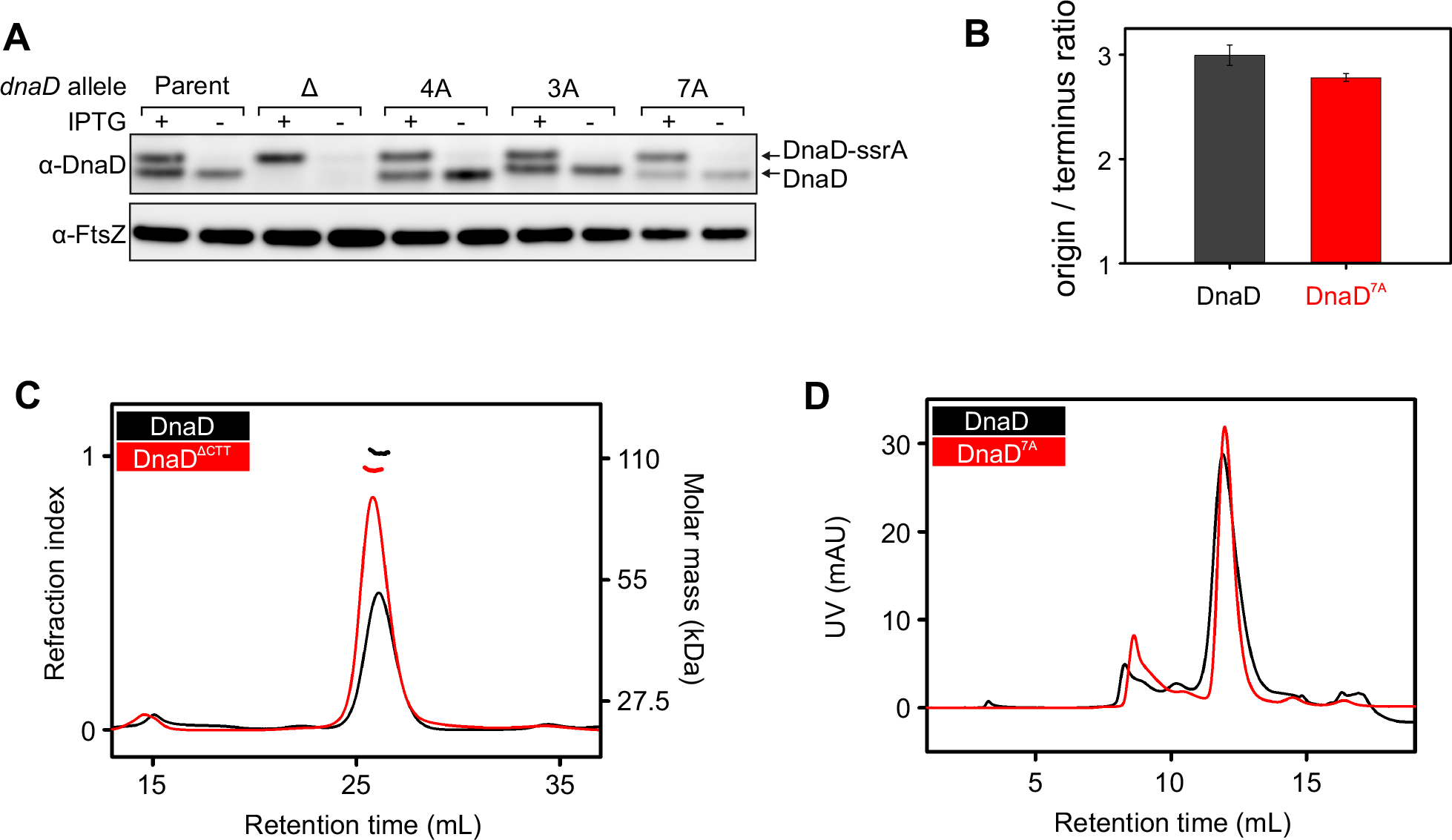
DnaD ssDNA binding characterisation. **(A)** Immunobloting of multiple alanine substitution DnaD variants targeting positively charged and aromatic residues within the C-terminal tail. The tubulin homolog FtsZ was used as a loading control. **(B)** Marker frequency analysis of the *dnaD^7A^* mutant using quantitative PCR. **(C)** SEC-MALS analysis of the DnaD variant lacking the C- terminal tail. **(D)** SEC analysis of DnaD^7A^.

**Figure S15.**
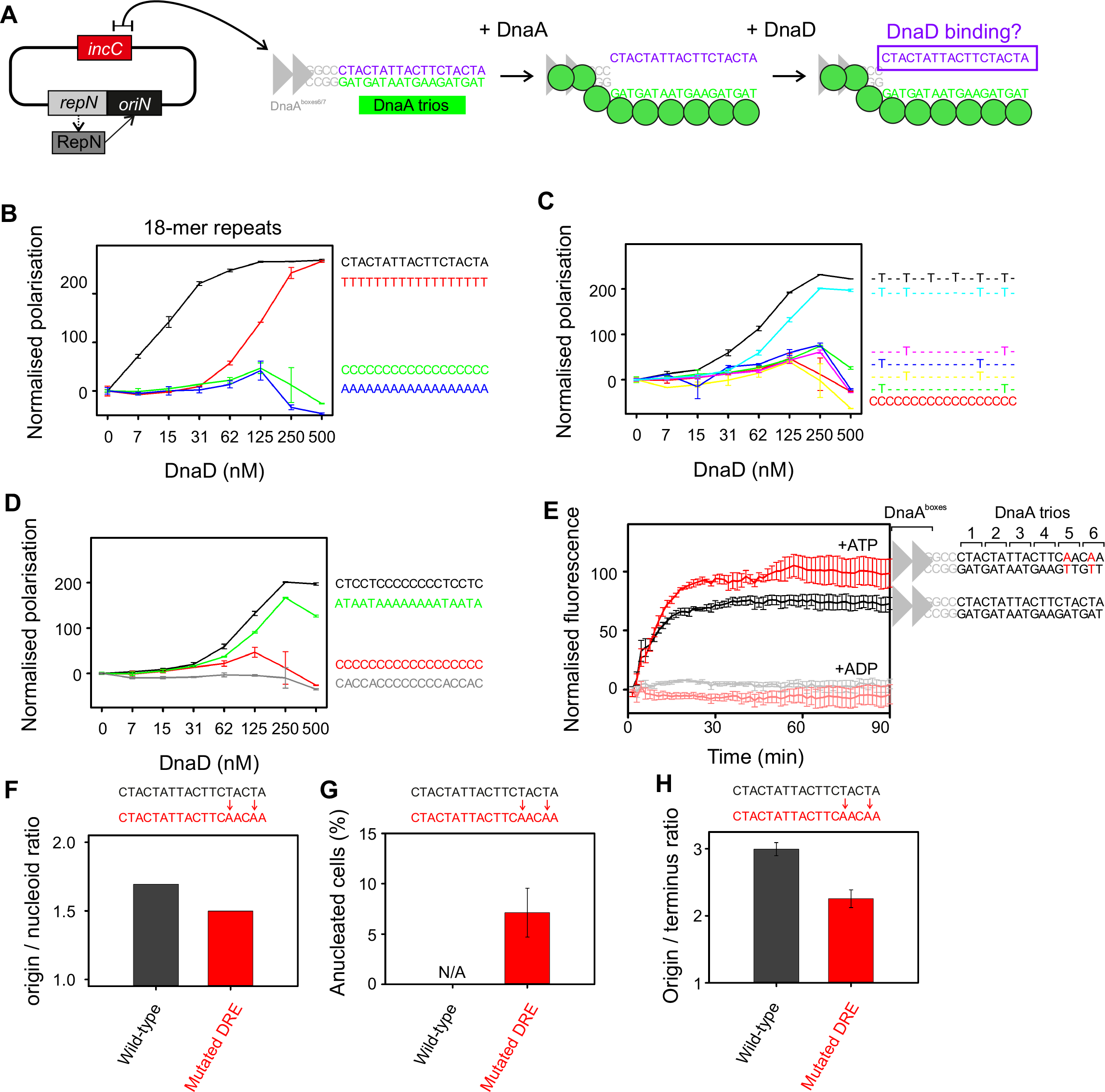
Two 5’-TnnT-3’ repeats are required for DnaD ssDNA binding *in vitro* and DNA replication initiation *in vivo*. **(A)** Illustration of the proposed basal origin unwinding mechanism involving DnaA oligomer formation on DnaA-trios. **(B)** Fluorescence polarisation analysis of DnaD binding homopolymeric 18-mers. **(C)** Fluorescence polarisation analysis of DnaD binding 5’-TnnT-3’ motifs located within an inert ssDNA substrate**. (D)** Fluorescence polarisation analysis of DnaD binding 5’-TnnT-3’ motifs located within an inert ssDNA substrate, and binding 5’-AnnA-3’ motifs. **(E)** Strand separation assay showing that mutating the distal 5’-TnnT-3’ element relative to the DnaA-boxes does not affect DnaA strand separation activity *in vitro*. **(F)** Quantification of origins per nucleoid in the DRE mutant background based on microscopy images taken following cell growth at 20°C. **(G)** Quantification of anucleated cells found in the DRE mutant background over the count of 750 cells from microscopy images taken following cell growth at 37°C. **(H)** Marker frequency analysis of the DRE mutant at 37°C measured using quantitative PCR.

**Figure S16.**
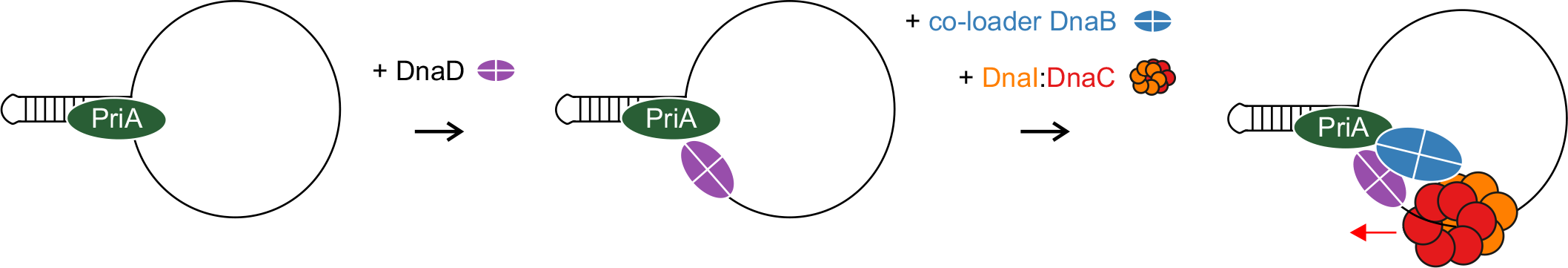
Model for helicase recruitment and loading in *B. subtilis* during PriA-dependent replication restart at a single-strand origin (*sso)*.

