## Supplement for "Positive and negative control of helicase recruitment at a bacterial chromosome origin"

**Supplementary Methods**

*Strains*

*B. subtilis* strains are listed in Supplementary Table 1. Transformation of competent *B. subtilis* cells was performed using an optimized two-step starvation procedure as previously described (Anagnostopoulos and Spizizen, 1961; Hamoen et al., 2002). Briefly, recipient strains were grown overnight at 37°C in transformation medium (Spizizen salts supplemented with 1 µg/ml Fe-$NH_4$-citrate, 6 mM $MgSO_4$, 0.5% glucose, 0.02 mg/ml tryptophan and 0.02% casein hydrolysate) supplemented with IPTG where required. Overnight cultures were diluted 1:17 into fresh transformation medium supplemented with IPTG where required and grown at 37°C for 3 hours with continual shaking. An equal volume of prewarmed starvation medium (Spizizen salts supplemented with 6 mM $MgSO_4$ and 0.5% glucose) was added and the culture was incubated at 37°C for 2 hours with continual shaking. DNA was added to 350 µl cells and the mixture was incubated at 37°C for 1 hour with continual shaking. 20-200 µl of each transformation was plated onto selective media supplemented with IPTG where required and incubated at 37°C for 24-48 hours. The genotype of all chromosomal *dnaA* and *dnaD* mutants was confirmed by DNA sequencing.

**CW197** (*trpC2* Δ*dnaD amyE*::*spc*($P_{HSA+1T}$-*dnaD-ssrA* $lacI^{Q18M/W220F}$) was constructed to study potentially lethal mutants of *dnaD in vivo* (Fig. 1B). First, an ectopic copy of *dnaD* was placed under the control of an IPTG-inducible promoter ($P_{HYPERSPANK}$)(Wagner et al., 2009). When combined with a deletion mutant of the native *dnaD*, the basal expression level of the ectopic copy was sufficient to sustain growth. Two approaches were taken to reduce the basal expression of the ectopic *dnaD*, lowering promoter activity and reducing DnaD stability. Promoter activity was inhibited by altering the transcription start site from A to T ($P_{HAT}$) and introducing mutations into *lacI* (Q18M/W220F) that increase operator binding (Daber and Lewis, 2009; Gatti-Lafranconi et al., 2013)(Fig. S2A). DnaD stability was reduced by fusing the ectopic *dnaD* to an *ssrA* degradation tag (AANDENYSENYALGG)(Griffith and

Grossman, 2008). Together these modifications produced a suitable expression system that conditionally complements the *dnaD* deletion mutant only when the ectopic *dnaD-ssrA* is induced (Fig. S2B). Immunoblot analysis confirmed nearly complete degradation of DnaD-ssrA following removal of IPTG within 30 minutes (Fig. S4C).

**CW198** (MDS42 *ΔrecA Δpdu Δrnh::kan*) was constructed by P1 transduction of *Δrnh*::*kan* from JW0204 (Keio collection) into CW181 (MDS42 *ΔrecA Δpdu* (Posfai et al., 2006)) harbouring pEAW365 (pET21A $P_{recA}$-*recA*), a gift from Elizabeth Wood and Michael Cox), followed by propagation in the absence of ampicillin to lose pEAW365.

**CW252** (*trpC2 amyE::spec(lacI* $P_{HYPERSPANK}$*-sirA) ganA::erm(xylR* $P_{XYL}$*-dnaD)*) was constructed by transformation with a PCR product generated by three-way Gibson assembly (NEBuilder HiFi). The *dnaD* gene with its native ribosome binding site was amplified using oCW611 and oCW612 using 168CA genomic DNA as template. The flanking region containing *ganA'-xylR-*$P_{XYL}$ was amplified using oCW284 and oCW614 with pJMP1 as template. The flanking region containing *erm-'ganA* was amplified using oCW613 and oCW130 with pJMP1 as template.

**CW270** (*trpC2 amyE::spec(lacI* $P_{HYPERSPANK}$*-sirA) ganA::erm(xylR* $P_{XYL}$*-*$dnaD^{I83A}$)) was constructed identically to CW252 with the exception of $dnaD^{F51A}$, $dnaD^{I83A}$ and $dnaD^{E95A}$ being amplified using oCW611 and oCW612 from CW174, CW170 and CW166 genomic DNA respectively as templates.

**HM1784** (BTH101 *Δrnh*::*kan*) was constructed by P1 transduction of *Δrnh*::*kan* from JW0204 (Keio collection) into BTH101 [F-, *cya-99*, *ara*D139, *gal*E15, *gal*K16, *rps*L1 ($Str^r$), *hsd*R2, *mcr*A1, *mcr*B1 (Euromedex)].

**HM1792** (DH5α *Δrnh*::*kan*) was constructed by P1 transduction of *Δrnh*::*kan* from JW0204 (Keio collection) into HM1785 (DH5α harbouring pEAW365 (pET21A $P_{recA}$-*recA*)), followed by propagation in the absence of ampicillin to lose pEAW365.

**DnaD alanine substitution strains** were generated by a blue/white screening assay using CW197 as parental strain and mutant plasmids obtained after Quickchange mutagenesis and sequencing as recombinant DNA (Fig. S3). X-gal 0.016% w/v was added to transformation plates for detection of β-galactosidase activity and selection of kanamycin resistant white colonies that integrated mutant DNA by double-recombination. Three individual white colonies per mutant were then restreaked onto a medium either with or without IPTG to identify alleles of interest.

*Plasmids*

Plasmids are listed in the Supplementary Table 2 (sequences are available upon request). DH5α [$F^-$ Φ80*lac*ZΔM15 Δ(*lac*ZYA-*arg*F) U169 *rec*A1 *end*A1 *hsd*R17($r_K^-$, $m_K^+$) *pho*A *sup*E44 *thi*-1 *gyr*A96 *rel*A1 $\lambda^-$] (Taylor et al., 1993) was used for plasmid construction, except for plasmids harbouring *dnaD* that were constructed in CW198. Descriptions, where necessary, are provided below.

**DnaD alanine-scan mutant plasmids** were generated by Quickchange mutagenesis using oligonucleotides listed in Supplementary Table 5. Cloning protocols were adapted to a 96-well plate format for PCR amplification of mutant plasmids, heat-shock and transformation recovery. All plasmids were sequenced.

**pCW123, pCW141, pCW142, pCW143, pCW153, pCW163, pCW214, pHM543, pHM544, pHM545** were generated by Quickchange mutagenesis using the oligonucleotides listed in Supplementary Table 3**.**

**pCW4** was generated by cloning *HindIII-SphI* PCR fragments generated using oligonucleotides listed in Supplementary Table 3.

**pDS84, pDS119, pDS120, pDS126, pDS127, pDS132, pHM359, pHM638, pHM640, pHM642, pHM644** were generated by cloning *Asp718I-BamHI* PCR fragments generated using the oligonucleotides listed in Supplementary Table 3**.**

**pCW66, pCW137, pCW171, pCW213, pSP075, pSP080, pSP081, pSP082, pSP083, pSP085** were generated by ligase-free cloning via two-step assembly processes using oligonucleotides listed in Supplementary Table 3. The underlined part of each primer indicates the region used to form an overlap. FastCloning (Li et al., 2011) was used with minor modifications. PCR products (15 µl from a 50 µl reaction) were mixed and then subjected to a heating/cooling regime: two cycles of 98°C for 2 minutes then 25°C for 2 minutes, then one cycle of 98°C for 2 minutes then 25°C for 60 minutes. After cooling *DpnI* restriction enzyme (1 µl) was added to digest parental plasmids and the mixtures were incubated at 37°C for ~4 hours. Following digestion 10 µl of the PCR mixture was transformed into chemically competent *E. coli*. Where several primer pairs are listed for the construction of a single plasmid (Multi-step assembly column in Supplementary Table 3), multiple rounds of ligase free cloning were performed to obtain the final constructs.

*Oligonucleotides*

All oligonucleotides were purchased from Eurogentec. Oligonucleotides used for plasmid construction are listed in Supplementary Table 3, oligonucleotides used for qPCR are listed in Supplementary Table 4 and those generated by the Quickchange program are listed in Supplementary Table 5.

**Quickchange mutagenesis** was used for the construction of the DnaD mutant plasmid library. Each point mutant was assembled by PCR using mutagenic primers carrying a single alanine substitution(Liu and Naismith, 2008). We generated all mutant primer pairs via an in-house Quickchange program that optimised sequences according to key features in site-directed mutagenesis primer design. These include sequence length adjustments based on: (i) the melting temperature ($T_M$) of the oligonucleotide part that anneals to the template plasmid, (ii) the $T_M$ corresponding to a primer pair overlapping section, (iii) the GC-content within different sections of individual primers, (iv) the presence of a GC-clamp at every oligonucleotide 3'-end, and (v) the $T_M$ difference between forward and reverse primer pairs. Code was written in Java and is available upon request.

*ChIP*

Strains were grown overnight at 30°C in Spizizen salts supplemented with tryptophan (20 µg/ml), glutamate (0.1%), glucose (0.5%) and casamino acid (0.2%). The following day cultures were diluted 1:100 into fresh medium and allowed to grow to an $A_{600}$ of 0.4. Samples were resuspended in PBS and cross-linked with formaldehyde (final concentration 1%) for 10 min at room temperature, then quenched with 0.1 M glycine. Cells were pelleted at 15°C, washed three times with PBS (pH 7.3) then frozen in liquid nitrogen and stored at -80°C. Frozen cell pellets were resuspended in 500 µl of lysis buffer (50 mM NaCl, 10 mM Tris-HCl pH 8.0, 20% sucrose, 10 mM EDTA, 100 µg/ml RNase A, ¼ complete mini protease inhibitor tablet (Roche), 2000 K u/µl Ready-Lyse lysozyme (Epicentre)) and incubated at 37°C for 30 min to degrade the cell wall. 500 µl of immunoprecipitation buffer (300 mM NaCl, 100 mM Tris-HCl pH 7.0, 2% Triton X-100, ¼ complete mini protease inhibitor tablet (Roche), 1 mM EDTA) was added to lyse the cells and the mixture was incubated at 37°C for a further 10 min before cooling on ice for 5 min. DNA samples were sonicated (40 amp) four times at 4°C to obtain an average fragment size of ~500 to 1000 base pairs. Cell debris were removed by centrifugation at 4°C and the supernatant transferred to a fresh Eppendorf tube. To determine the relative amount of DNA immunoprecipitated compared to the total amount of DNA, 100 µl of supernatant was removed, treated with Pronase (0.5 mg/ml) for 60 min at 37°C then stored on ice.To immunoprecipitate protein-DNA complexes, 800 µl of the remaining supernatant was incubated with rabbit polyclonal anti-DnaA, anti-DnaD and anti-DnaB antibodies (Eurogentec) for 1 hour at room temperature. Protein-G Dynabeads (750 µg, Invitrogen) were equilibrated by washing with bead buffer (100 mM $Na_3PO_4$, 0.01% Tween 20), resuspended in 50 µl of bead buffer, and then incubated with the sample supernatant for 1 hr at room temperature. The immunoprecipitated complexes were collected by applying the mixture to a magnet and washed once with the following buffers for 15 min in the respective order: 0.5X immunoprecipitation buffer; 0.5X immunoprecipitation buffer + NaCl (500 mM); stringent wash buffer (250 mM LiCl, 10 mM Tris-HCl pH 8.0, 0.5% Tergitol-type NP-40, 0.5%

sodium deoxycholate 10 mM EDTA). Finally, protein-DNA complexes were washed a further three times with TET buffer (10 mM Tris-HCl pH 8.0, 1 mM EDTA, 0.01% Tween 20) and resuspended in 100 µl of TE buffer (10 mM Tris-HCl pH 8.0, 1 mM EDTA). Formaldehyde crosslinks of both the immunoprecipate and total DNA was reversed by incubation at 65°C for 16 hours in the presence of 1,000 U Proteinase K (excess). The reversed DNA was then removed from the magnetic beads, cleaned using QIAquick PCR Purification columns (Qiagen) and used for qPCR analysis.

*qPCR*

To measure the amount of genomic loci bound to DnaA, DnaD and DnaB, the Luna qPCR mix (NEB) was used for PCR reactions and qPCR was performed in a Rotor-Gene Q Instrument (Qiagen) using serial dilutions of the immunoprecipitate and total DNA control as template. Oligonucleotide primers were designed to amplify *oriC* (qSF11/qSF12), *oriN* (qSF5/qSF6) and the non-specific locus *yhaX* (oWKS145/oWKS146 (Smits et al., 2011)), and were typically 20–25 bases in length and amplified a ~100 bp PCR product (Supplementary Table 4). Error bars indicate the standard error of the mean for 6-8 biological replicates.
